# Targeted sequencing of FH-deficient uterine leiomyomas reveals biallelic inactivating somatic fumarase variants and allows characterization of missense variants

**DOI:** 10.1101/663609

**Authors:** Bernt Popp, Ramona Erber, Cornelia Kraus, Georgia Vasileiou, Juliane Hoyer, Stefanie Burghaus, Arndt Hartmann, Matthias W. Beckmann, André Reis, Abbas Agaimy

**Affiliations:** Institute of Human Genetics, University Hospital Erlangen, Friedrich-Alexander-Universität Erlangen-Nürnberg (FAU), Erlangen, Germany; Institute of Human Genetics, University of Leipzig Hospitals and Clinics, Leipzig, Germany; Institute of Pathology, University Hospital Erlangen, Friedrich-Alexander-Universität Erlangen-Nürnberg (FAU), Erlangen, Germany; Department of Obstetrics and Gynecology, University Hospital Erlangen, Comprehensive Cancer Center ER-EMN, Friedrich-Alexander University Erlangen-Nürnberg (FAU), Erlangen, Germany

**Keywords:** uterine leiomyoma, fumarate hydratase, fumarase, FH, HLRCC, panel sequencing, missense

## Abstract

Uterine leiomyomas (ULs) constitute a considerable health burden in the general female population. The fumarate hydratase (FH) deficient subtype is found in up to 1.6% and can occur in hereditary leiomyomatosis and renal cell carcinoma (HLRCC) syndrome.

We sequenced 13 FH deficient ULs from a previous immunohistochemical screen using a targeted panel and identified biallelic *FH* variants in all. In eight, we found a *FH* point mutation (two truncating, six missense) with evidence for loss of the second allele. Variant allele-frequencies in all cases with a point mutation pointed to somatic variants. Spatial clustering of the identified missense variants in the lyase domain indicated altered fumarase oligomerization with subsequent degradation as explanation for the observed FH deficiency. Biallelic *FH* deletions in five tumors confirm the importance of copy number loss as mutational mechanism.

By curating all pathogenic *FH* variants and calculating their population frequency, we estimate a carrier frequency of up to 1/2,563. Comparing with the prevalence of FH deficient ULs, we conclude that most are sporadic and estimate 2.7 - 13.9% of females with an FH deficient UL to carry a germline *FH* variant.

Further prospective tumor/normal sequencing studies are needed to develop a reliable screening strategy for HLRCC in women with ULs.

## INTRODUCTION

Uterine leiomyomas (ULs; fibroids) are benign smooth muscle tumors of the myometrium with an estimated lifetime risk of 70%(1). As about 30% of women are symptomatic and present with abdominal pain, vaginal bleeding or anemia(2, 3), ULs represent a considerable health burden(4). These hormone dependent tumors usually do not occur before adolescence, but increase in size in the reproductive period and frequently decrease in size after menopause(2). Besides these hormonal influences other factors associated with modulating the individual risk for ULs are certain dietary habits, caffeine and alcohol consumption, smoking, components of the metabolic syndrome (central obesity, high blood pressure, hyperlipidemia) and ethnic background(5). The observation that ULs are more common in women of African origin than in Caucasian women(6) implicates genetic factors as additional risk factors.

Genome-wide association studies (GWAS) have linked several genomic regions and biological processes like mRNA degradation(7), thyroid function(8) and fatty acid synthesis(9) with UL risk. A large study(10) identified several novel loci associated with estrogen metabolism, uterine development and genetic instability. The authors could associate the combined polygenic risk with the most common UL subtype, positive for somatic *MED12* mutations(10).

From a molecular pathologist view, there are currently four different and mutually exclusive groups of ULs defined by their typical driver variants(11). ULs with either *MED12* hotspot driver mutations or with genomic rearrangements involving *HMGA2* are the most common and represent up to 90%(12). The two other known subtypes, defined by typical deletions in the *COL4A5*/*COL4A6* genes or *FH* deficiency, are much rarer. The *COL4A5*/*COL4A6* deletion positive UL subtype constitutes about 2%(12). We and other estimated that FH deficient UL subtype makes up 0.4% to 1.6% of all ULs(13, 14).

Both rare subtypes are associated with certain highly penetrant heritable mendelian syndromes. Recurrent deletions at the *COL4A5*/*COL4A6* gene locus lead to the X-linked dominantly inherited diffuse leiomyomatosis with Alport syndrome (DL-ATS; OMIM #308940). Germline Loss-of-Function (LOF) variants (both truncating and missense) in the fumarate hydratase (fumarase) gene *FH* cause the dominantly inherited hereditary leiomyomatosis and renal cell cancer syndrome (HLRCC; OMIM #150800). The latter is characterized by multiple cutaneous and uterine leiomyomata and increased risk for aggressive renal cell carcinomas. While both monogenic heritable UL subtypes are rare, their occurrence in a syndromic setting with additional health problems in affected individuals and increased risk for relatives makes a timely diagnosis particularly important.

FH deficient ULs can be suspected based on morphological classification (“FH-d morphology”) divided in 1) low and 2) high magnification features(13-21). These include 1a) prominent branching blood vessels with thin walls (also: “staghorn” or “hemangiopericytoma-like” vessels)(19), 1b) a certain edema pattern (“alveolar”)(16), 1c) rhythmic neurilemoma-like (“chain-like”) arrangement of nuclei(14), 1d) bizarre nuclei(16, 22), 2a) eosinophilic cytoplasmic inclusions(17) and 2b) enlarged nuclei (“macronucleoli”) with perinucleolar halos(21). Despite the broad adaption of the FH-d morphology, its sensitivity/specificity and reproducibility are unproven. Immunohistochemistry (IHC) can be used to confirm the morphological suspicion. Loss of staining for the *FH* gene product fumarate hydratase is an intuitive direct marker, but recent reports of retained staining in tumors with pathogenic missense variants raised concerns about its validity as marker for FH deficiency(15, 18). In contrast, staining for 2SC (S-(2-succinyl) cysteine), a covalent protein modification that accumulates when the fumarate hydratase in cells is non-functional, has excellent statistical performance despite being an indirect marker of enzyme function(19, 23, 24). Unfortunately, the 2SC antibody is currently not commercially available which excludes the use in a clinical setting(15, 16, 18). Despite considerable research efforts, the question about an optimal pathologist-based screening method for HLRCC-associated ULs remains unanswered.

While ULs are common and benign tumors, uterine leiomyosarcomas (ULMSs) are rare malignant tumors of the myometrium. Recent reports have raised the hypothesis that ULMSs can rarely originate from pre-existing ULs(25, 26). The TCGA study has identified frequent monoallelic (78%) and often biallelic (14%) deletions of the tumor suppressor gene *RB1* as a driver in ULMSs(27). Monoallelic *RB1*-loss has been described in few ULs(25, 28), raising the questions 1) if it is a pre-existing event in certain ULs and a second-hit can induce malignancy and 2) whether routine RB1 screening, for example by IHC, is useful to identify high risk ULs.

Based on our cohort previously screened on morphological features and by FH IHC, we now analyzed 13 FH deficient ULs with available tumor DNA by targeted panel sequencing to investigate their somatic variant spectrum for both small genetic (single nucleotide variants: SNVs, base insertions or deletions: indels) and copy number variants (CNVs) and additionally characterized these ULs by RB1 IHC.

## MATERIALS AND METHODS

### Included samples and individuals

We included 13 FH deficient ULs with sufficient high-quality DNA from a previously described cohort of 22 prospectively diagnosed tumors collected from routine surgical pathology (n = 10) or consultation files (n = 3) of one of the authors (A.A.)(13). The histopathological characteristics of theses ULs have been reviewed by an experienced pathologist. Genetic counseling and molecular genetic testing were recommended if FH loss was identified in IHC routine. Detailed cohort descriptions are provided in Supplementary data file 1 sheet “cohort”.

### Immunohistochemistry

FH immunohistochemistry (IHC) had been performed and the method described previously(13). In brief, sections from formalin-fixed paraffin embedded (FFPE) tumor blocks were stained on a Ventana BenchMark Ultra automated instrument using the ultraView Universal DAB Detection Kit (both: Ventana Medical Systems, Inc., Tucson, USA) according to the institutes routine standards. Heat-induced epitope retrieval (HIER) was performed using cell conditioning solution 1 (CC1; Ventana Medical Systems, Inc., Tucson, USA) at 95 °C for 36 min followed by antibody incubation at 37 °C for 32 min using the mouse monoclonal antibody clone J-13 (Santa Cruz Biotechnology, Inc., Dallas, USA) at 1:50 dilution. FH expression was classified into four categories (intact, loss/deficient, aberrant expression, not assessable). Aberrant expression as defined as any abnormal looking pattern other than the three patterns listed above.

RB1 IHC was performed on 12 of the 13 ULs with a mouse monoclonal antibody (Clone G3-245, 1:100, BD Biosciences Pharmingen, Franklin Lakes, USA) according to the same procedures described for FH IHC. RB1 expression was classified into five categories. Intact expression was defined as tumor and internal control (e.g. endothelial cells) with strong nuclear positive staining. Reduced expression was defined as weaker staining intensity of tumor nuclei compared to the internal control. Loss/deficiency was defined as completely negative tumor nuclei with intact internal control. A hybrid pattern was defined as areas with loss next to areas/cells with intact expression. When both tumor nuclei and internal control were negative the case was categorized as not assessable.

IHC classification for both FH and RB1 was performed according to above classifications by two independent observers (R.E., A.A.) blinded to clinical data.

### Panel sequencing of UL tumor DNA

Genomic DNA was extracted from formalin-fixed paraffin-embedded (FFPE) tumor tissue using the QIAamp DNA FFPE Tissue kit (QIAGEN, Redwood City, USA) after areas with tumor cellularity of >80% were marked by a trained pathologist and manually macro-dissected. Enrichment and library preparation were performed with the TruSight Cancer Sequencing Panel v1 (Illumina, Inc., San Diego, USA) using 500ng of genomic DNA. This commercial panel includes 95 genes associated with different cancer syndromes (Supplementary file 1 sheet “trusight_cancer_v1_genes”). Besides the higher DNA input, all procedures were performed according to the manufacturers’ instructions. Libraries were sequenced with 150 bp paired end reads on an MiSeq system (Illumina, Inc., San Diego, USA).

After demultiplexing, quality and adapter trimming was performed on the reads using cutadapt(29) version 1.10 from within the wrapper tool Trim Galore version 0.4.1. Read alignment to the hg19 reference genome was performed with BWA-MEM(30) version 0.7.15. After removing duplicate reads with Picard tools version 2.5.0 and local realignment of indels, tumor-only variant calling was performed on the final tumor BAM files (for alignment statistics see Supplementary file 1 sheet “panel_stats”) using MuTect2(31) from GATK version 3.7-0(32) and a panel of 100 in-house germline control samples sequenced on the same platform. For annotation of the resulting variant files, SnpEff and SnpSift(33, 34) were used with dbNSFP(35) version 2.93 and variant frequencies from the ExAC database(36) version 3.1 and COSMIC database(37) version 79 based on the files provided from the respective website. The annotated variants were filtered to have a coverage (DP) of at least 10 reads and an allele fraction (AF) of at least 10%. Variants present in the ExAC(36) database ≥ 100 times were filtered out unless they were also reported in the COSMIC database ≥ 10 times or were reported as (likely) pathogenic in ClinVar(38). Only coding and splice site variants were further analyzed. Subsequently, the resulting lists were examined using the IGV browser(39) and evaluated for their biological plausibility.

CNV calling from panel data was performed on the same BAM-files used for variant calling utilizing CNVkit(40) version 0.8.3 with standard parameters against the same 100 germline control samples used for variant calling. The CNVkit “call” command was used with a purity setting of 0.8 to convert log2 ratios into integer CN-(copy number) values. Results were visualized with the “scatter” and “heatmap” functions in CNVkit.

### Estimation of germline probability

To estimate whether the *FH* SNVs/indels identified from our tumor-only sequencing approach were germline or somatic variants, we first generated a plausible range of purity estimates for uterine tumors from published reports. As we could not identify large studies estimating the purity of macro-dissected ULs from sequencing data, we used purity estimates(41) of two other uterine tumor types (uterine carcinosarcoma, uterine endometrial carcinoma) from The Cancer Genome Atlas Program (TCGA)(42). We then plotted the theoretical relationship between tumor purity (TP) and expected variant allele fraction (VAF) for germline and somatic variants (formula: “AF = TP + (1 - TP) * initial zygosity in the germline”) assuming a somatic second-hit deleting the other allele and compared this to the variant allele frequency observed in our samples (see Figure S3 for methodological details).

### Collection and computational analyses of *FH* variants

To analyze enrichment of likely disease associated *FH* variants in domains, we downloaded all described variants from the ClinVar(38), LOVD(43, 44) and COSMIC(37) databases and scored these with InterVar(45) according to the American College of Medical Genetics and Genomics (ACMG) 5-tier classification(46), which is typically used to assess the pathogenicity of SNVs/indels in a clinical setting.

To assess the population carrier frequency, we annotated all *FH* variants from the public population databases gnomAD (https://gnomad.broadinstitute.org/) and BRAVO (https://bravo.sph.umich.edu/) with our curated set of (likely) pathogenic variants (see Table S2, Table S3 and Supplementary file 2 for details).

By generating all missense variants possible though a single base exchange for *FH* and plotting different computational scores, along the linear protein representation, we analyzed domain regions intolerant to missense variation. CADD(47), M-CAP(48) and REVEL(49) are recently developed scores utilizing different information types like conservation and functional consequences. They are typically used to access the pathogenicity of a missense variant.

All variant-sets were harmonized to a common reference with VariantValidator(50) (NM_000143.3 transcript, hg19 reference genome) and annotated with the same pipeline (Supplementary notes) to guarantee a uniform naming.

### Protein structure analysis of identified *FH* missense variants

We visualized the spatial clustering of identified likely somatic *FH* missense variants in 3D using the publicly available tertiary protein structure data of human fumarase (PDB-ID: 5D6B)(51) with the Pymol molecular visualization software (Version 1.8.6.0; Schrödinger LLC, New York, USA) installed through Conda (Anaconda Inc., Austin, USA).

To estimate the probability of the observed spatial clustering, we employed the online version of mutation3D(52), which uses a bootstrapping approach to estimate an empiric p-value, with the 5D6B protein structure as template. As a baseline, we compared the 3D clustering analysis of the herein identified six likely somatic *FH* missense variants to all (likely) pathogenic missense variants from the curated ClinVar and LOVD datasets (Figure S2).

## RESULTS

### Study cohort

The median age at diagnosis in the 13 females included was 37 years (y) with a range of 25y to 72y. Seven individuals were treated by hysterectomy and five by enucleation (no data for one case). Eight individuals had more than one UL nodule (no data for one case). Three individuals had a personal history of tumors/cancer of other organ systems including thyroid adenoma (S06), colorectal adenocarcinoma and endometrioid carcinoma of the uterus (S07) and breast cancer (S12). For three individuals, a family history of tumors/cancer in first degree relatives was reported to the pathologist: lung carcinoma at age 56y in the mother of S05, chronic leukemia at age 56y in the father and adenoma of the thyroid at age 30y in the mother of S06 and colorectal adenocarcinoma at age 69y in the sister and gastric carcinoma at age 84y in the father of S07. None of the 13 individuals presented with other typical features (cutaneous leiomyoma or renal tumors) of HLRCC besides their FH deficient ULs, indicating sporadic events.

Only one individual (S11) presented for genetic consultation at our Center for Rare Diseases. Molecular genetic testing identified no pathogenic germline variant in the *FH* gene. Clinical examination and family history raised the suspicion of tuberous sclerosis complex which could be confirmed by extended panel analysis and RNA analyses (detailed clinicopathological description of this case is provided in the Supplementary notes and Figures S7 and S8).

### FH loss and additionally reduced RB1 expression in IHC

All 13 ULs, selected for panel sequencing in this study, showed a complete loss of FH staining by IHC as reported previously(13). RB1 IHC showed a reduced expression in all 12 (100%) analyzed FH deficient ULs with no sample showing a complete loss. For one sample (S02), no RB1 IHC was performed.

### Properties of identified *FH* SNV/indels

Tumor-only variant calling for small genetic variants using a somatic variant caller identified a single SNV/indel in the *FH* gene in 8/13 (61.5%) of ULs. In no sample, we identified two SNVs/indels. Two of the identified variants (25.0%) were annotated as likely gene disrupting. The variant c.457delG in individual S06 causes a frame-shift which directly introduces a termination codon (p.(Val153*)). The variant c.379-2A>G in S03 disrupts the conserved splice-acceptor of exon 4, is predicted to cause aberrant splicing (denoted as “r.spl?” according to Human Genome Variation Society (HGVS) recommendations) by all five different computational splice prediction algorithms (Supplementary file 1 sheet “FH_SNVindel_summary”) and has been described to reduce FH activity(53).

The remaining six variants were annotated as missense variants causing the substitution of different single amino acids. Five of the missense variants were predicted as disrupting protein function throughout all eight computational missense prediction scores used. Only the variant c.1236G>A, p.(Met412Ile) identified in S09 was predicted as benign by five out of eight scores. Instead, this variant is predicted to cause aberrant splicing by four of the five splice prediction tools used (Supplementary file 1 sheet “FH_SNVindel_summary”). The c.1236G>A variant changes the last coding guanin base of exon 8 into an adenine. This base position is typically highly conserved in the consensus splice-site sequence(54). Furthermore, c.1236G>A is the only variant annotated as missense which does not affect a conserved protein domain like the Lyase domain. This domain contains the other five missense variants and most of the (likely) pathogenic missense variants reported in databases (Figure 1A). Therefore, it seems likely that the c.1236G>A variant in fact disrupts normal splicing (HGVS nomenclature “r.spl?”) and should be regarded as a likely gene disrupting variant (HGVS nomenclature “p.0?”). Further studies on RNA are needed to confirm the exact consequence of this variant. For the other five missense variants, computational splice-effect prediction scores were either unremarkable or not available, which points to the conclusion that these are true missense variants.

**Figure 1.**
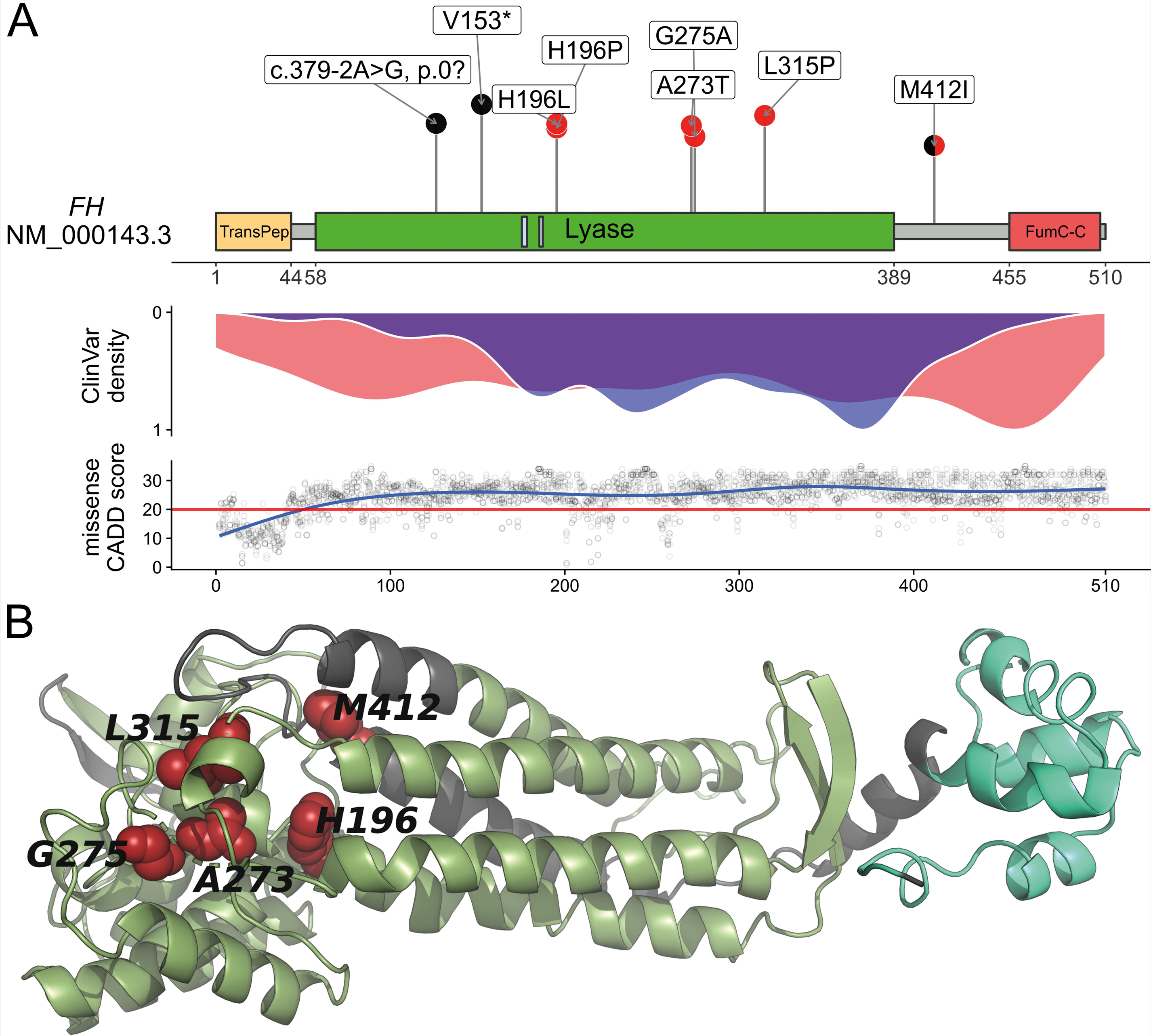
Somatic SNV/indel spectrum variant properties. (A) Upper panel: Schematic representation of the FH protein, domains (ticks below x-axis numbered after NP_000134.2 and P07954) and localization of herein identified likely somatic variants. Likely gene disrupting variants are presented in black and missense variants in red. TransPep, mitochondrion transit peptide; Lyase, N-terminal fumarate lyase domain; small light blue boxes in Lyase domain: substrate binding site A and B; FumC-C, C-terminal fumarase C domain. Middle panel: Density plot showing the distribution of (likely) pathogenic truncating (red) and missense (blue) variants reported in the publicly available database ClinVar(38) which collects user submitted curations for variant pathogenicity. Lower panel: Generalized linear models of the CADD(47) score, a computational (“in silico”) metric commonly used to assess the possible pathogenicity of single nucleotide variants based on diverse annotations, for all possible *FH* missense variants. (B) Structural analysis of human fumarase (FH, dark grey) based on the protein crystal structure with PDB (Protein Data Bank; https://www.rcsb.org) code 5D6B(51). Green, Lyase domain; pale green, FumC-C domain; red, amino acid residues for the mutations L315P, H196P, G275A, M412I, A273T, H196L lying in the Lyase domain or close to it. The missense mutations in the Lyase domain affect highly conserved amino acid residues which likely disrupt the protein structure (see also Figure S01 and S02). One letter amino acid code was used due to space constraint.

As missense variants are not expected to cause protein loss, but all 13 ULs were preselected based on complete FH-loss in IHC, the identification of true missense variants in 5/13 (38.5%) of ULs was unanticipated. When analyzing the proximity of the missense variants in the tertiary fumarase protein structure, we observed that the variants lie very close to each other (Figure 1B). We therefore performed 3D clustering analyses using mutation3D(52), which showed that the three variants identified in samples S01 (c.944T>C, p.(Leu315Pro)), S07 (c.824G>C, p.(Gly275Ala)) and S10 (c.817G>A, p.(Ala273Thr)) form a protein-wide significant cluster (p-value: 0.0411, empirical bootstrapping approach) within the fumarase protein (Figure S2).

While the genomic position of seven SNV/indel variants showed high read coverage allowing reliable estimation of VAF, only the variant c.824G>C, p.(Gly275Ala) in individual S07 was covered relatively low with 19 reads. All eight variants showed high VAF between 0.443 and 0.793 (median 0.766), indicating a CN-loss of the second allele as second-hit.

### Somatic variant status in FH deficient ULs

As SNV/indel calling only identified one *FH*-variant in eight of the 13 ULs, we next performed CN-calling from the capture-based sequencing data using the CNVkit algorithm(40) to search for second-hits. Assuming an average tumor purity of 80%, we were able to identify a CN-loss in all 13 ULs studied. The *FH* CN-losses ranged in size between 15.9 and 42,757.4 kilo-bases (kb) with a median of 4,897.1 kb (Figure 2A; Table 1; Supplementary file 2 sheet “FH_CNVkit_summary”). In the five ULs with no SNV/indel identified, the log2 ratios and CN-values indicate biallelic deletions. Due to the relatively low resolution of the panel (when compared to chromosomal microarrays), the likely independent two deletions in theses sample have often been called with the same breakpoints. In sample S12, the CNVkit algorithm indeed called a small 15.9 kb deletion affecting exons 3 to 10 of the *FH* gene and a large 1407.7 kb deletion in the chromosomal band 1q43 affecting the whole *FH* gene (Figure 2B and C). For seven of the eight ULs with a SNV/indel variant identified, the CN-analysis indicated the deletion of a single allele, confirming the suspicion from the observed high VAF. In UL sample S05, with the c.587A>T, p.(His196Leu) variant at a relatively low VAF of 0.443, a 15.9 kb small deletion was called. This deletion had an exceptionally low (more negative than the monoallelic deletions and less negative than the biallelic deletions) log2 value of - 1.87 when compared to the other seven samples with one deletion only (log2 value range - 0.77 to −1.22). Together with the low VAF of the SNV in this sample, this observation points to the explanation of tumor-heterogeneity in this UL with multiple second-hit events occurring after an initial deletion of one *FH* allele.

**Table 1.**
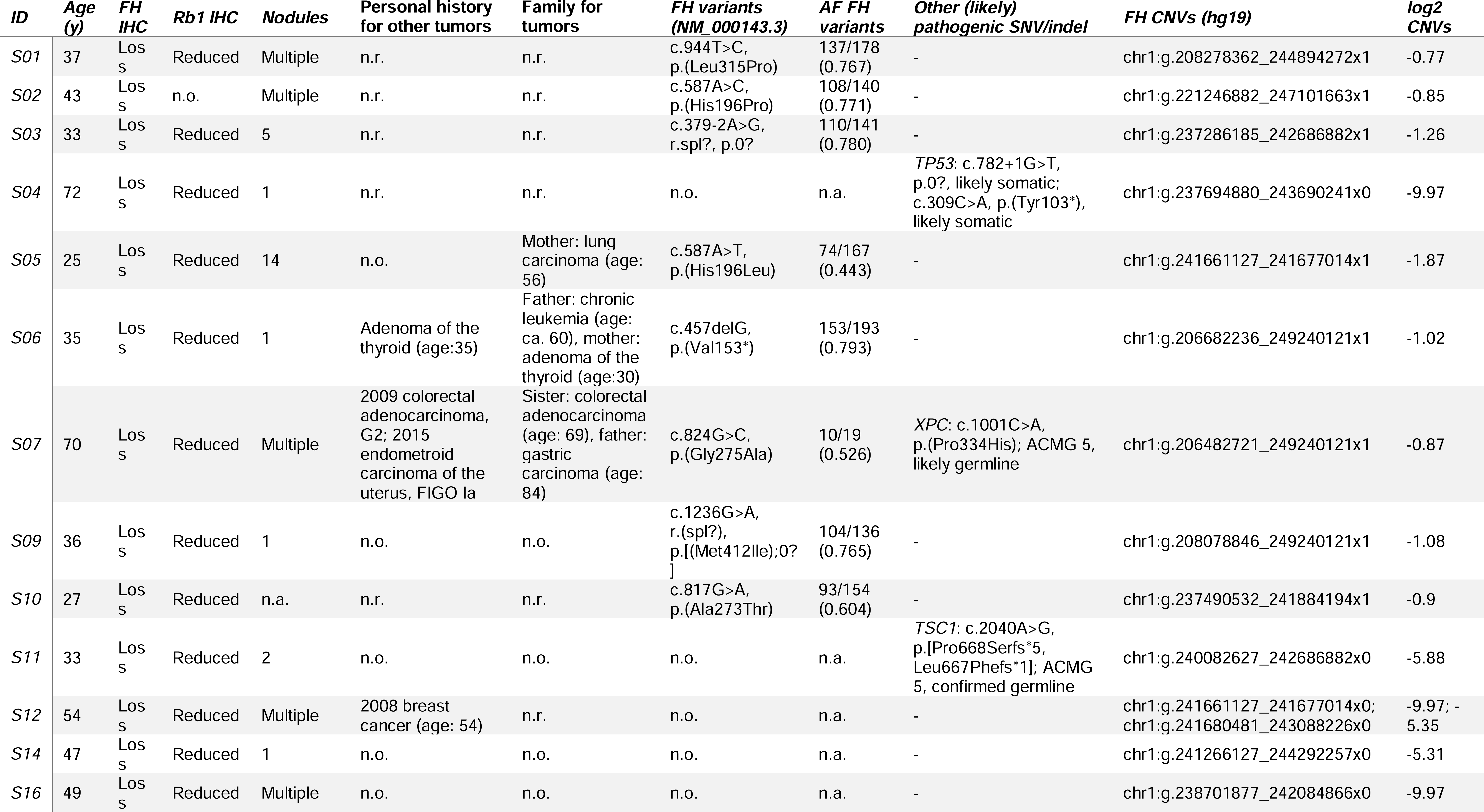
Overview of analyzed cases and identified somatic variant status. n.o. = none observed; n.r. = no reported; n.a. = not applicable; IHC = Immunohistochemistry; AF = allele fraction

**Figure 2.**
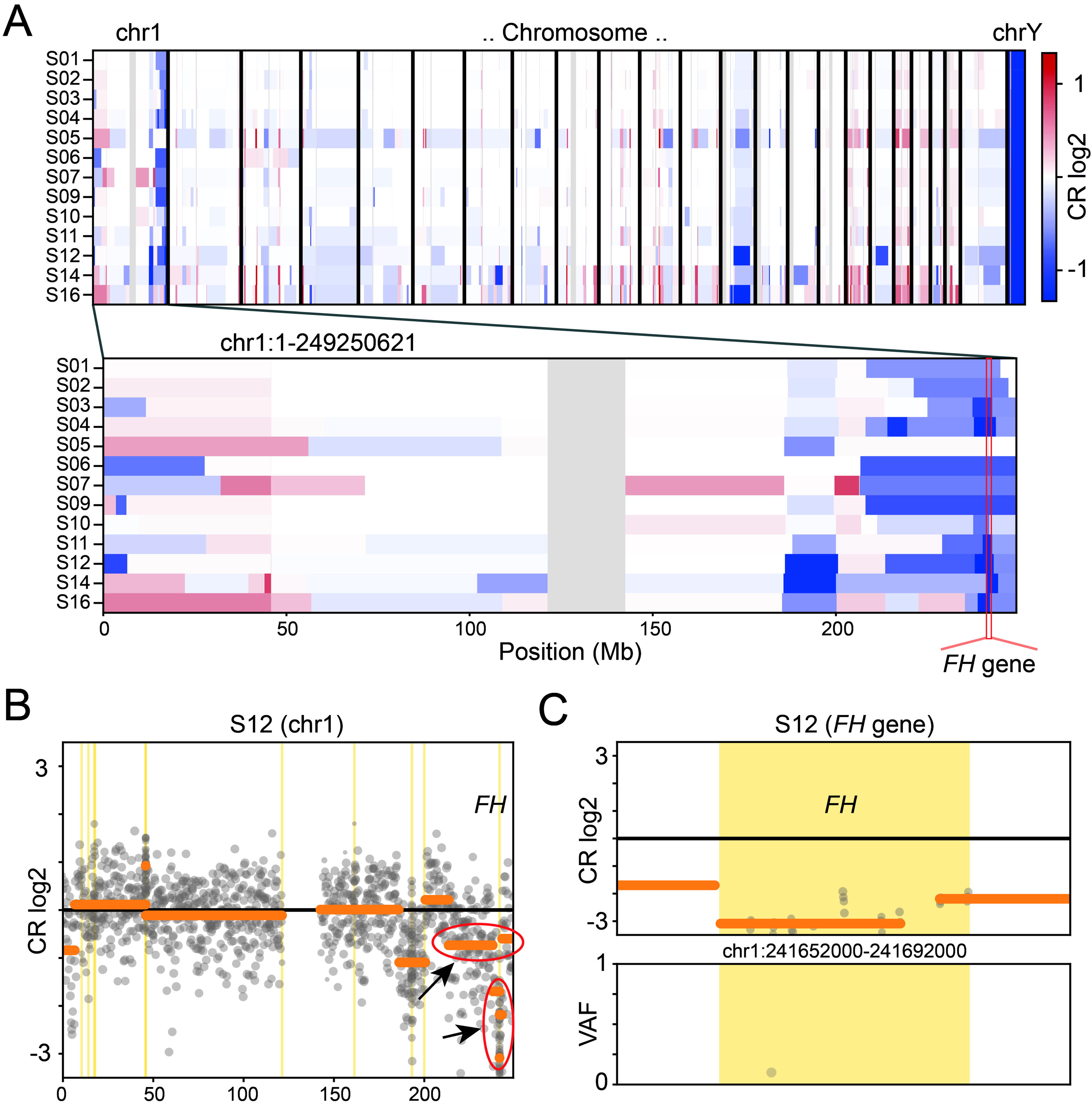
Somatic copy number aberrations. (A) Heat-plot of copy number (CN) aberrations detected in the 13 analyzed ULs using panel data. Upper panel: all chromosomes. Lower panel: zoomed in chromosome 1 containing the *FH* gene. Blue: CN loss, red: CN gain, CR: copy ratio. The position of the *FH* gene is indicated by the red stroke in the lower panel. Note the dark blue squares at the locus indicating samples with biallelic CN-loss in the tumor. In contrast to the relatively stable situation for SNVs/indels, all ULs show several larger CNVs with different recurrently affected genomic loci. When comparing the CNV-spectrum of the FH deficient ULs to 15 in-house HBOC tumors and correcting for multiple testing, only the *FH*-gene remained significant for CN-losses and the *ALK*-gene for CN-gains (see also Figure S4). Note that the 15.9 kb small deletion on one allele of sample S05 is merely identifiable at this resolution (compare Figure S5 CR-profile of this sample at the *FH*-gene locus). (B) Exemplary CR profile for sample S12 at chromosome 1 showing different rearrangements and especially the *FH*-gene locus affected by both a larger CN-loss and a second smaller one (marked by red ellipses and black arrows). (C) Zoomed in CR profile for sample S12 at the *FH*-gene locus with VAF (variant allele frequency) plot (lower panel). Grey dots represent markers used by CNVkit(40) (target and anti-target regions) and shading the dots indicates weight within the analysis. Vertical yellow bars mark gene regions. Horizontal orange bars represent CN segmentation calls.

The analysis of other genes covered by the panel identified the two pathogenic likely gene disrupting variants c.782+1G>T, p.0? and c.309C>A, p.(Tyr103*) in the *TP53* gene in UL sample S04 with VAFs (0.408 and 0.370, respectively) pointing to somatic variants. The VAFs of these two reliable somatic variants confirm our purity estimate of about 80% (e.g. multiply VAF by two for heterozygous cells: 0.408*2 = 0.816 and 0.370*2 = 0.740). No further likely somatic SNVs/indels were identified in the other 12 samples in all 95 genes covered by the panel. The only other pathogenic variant identified was the c.1001C>A, p.(Pro334His) in the *XPC* gene in UL sample S07, which is a known pathogenic variant (dbSNP rs74737358) causing the recessively inherited xeroderma pigmentosum (OMIM #278720) when inherited with a second pathogenic variant on the other allele. The high allele frequency of this variant in the general population (854/278,442 ∼ 0.31% in gnomAD v2.1.1) together with its VAF of 0.560 indicated that this is very likely a germline variant.

In contrast to the relatively stable situation for somatic SNVs/indels, all ULs showed several CN-aberrations especially on chromosome 1 (Figure 2A). Unsurprisingly, the *FH*-gene showed significantly more CN-losses in the 13 ULs, but no other gene reached panel-wide significance when correcting for multiple testing. Interestingly, the *RB1* gene locus indicated a deletion ≥ 500 kb in only three FH deficient ULs (two additional when considering CNVs < 500 kb), despite the reduced expression in IHC in all 12 samples analyzed. In regards of CN-gains, we found a panel wide significance only for the *ALK*-gene locus (Figure S4).

### Observed variant allele fractions are best explained by a somatic two-hit model

Assuming the typically relatively high purity of a usually monoclonal and non-invasive tumor like ULs, the VAF observed in eight ULs with an SNV/indel (between 0.443 and 0.793; see Table 1) are best explained by a somatic two hit model with an somatic SNV/indel on one allele and a second somatic CN-loss on the other allele (Figure S3). The alternative hypothesis of an initial germline variant would require an unusually low tumor purity (< 57.7%) to explain the observed VAF values herein.

### Carrier frequency for pathogenic *FH* variants in databases and prevalence of FH deficient ULs

By collecting described *FH* variants from the ClinVar and LOVD databases and classifying them according to ACMG criteria, we summarized 280 unique (likely) pathogenic variants. These included 130 protein truncating variants (46.4%), 36 variants affecting the splice regions (12.9%), 104 missense variants (37.1%) and 10 in-frame indel (3.6%) variants (Figure S1, Supplementary file 2). Truncating variants were distributed throughout the protein, while missense variants were enriched in the Lyase domain (Figure 1A middle panel, Figure S1).

By summarizing these 280 curated (likely) pathogenic *FH* variants in public databases, we estimated a carrier frequency (CF) of 1/2,563 (0.0390%) to 1/3,247 (0.0308%) individuals for the BRAVO and gnomAD population databases, respectively (Table S2, Table S3 and Supplementary file 2).

We calculated the expected prevalence of FH deficient ULs (prevFH-dUL) using published UL prevalence in the female population (prevUL) of up to 70% (1) and the proportion of FH deficient ULs (propFH-dUL) observed on pathologist routine(13, 14) between 0.3% and 1.1% (formula: propFH-dUL = prevUL * propFH-dUL = 70% * [0.4% to 1.6%] ≈ 0.3% to 1.1%).

Assuming independence between prevalence of FH deficient ULs and carrier frequency, only about 1/36 (calculation lower bound: 1 / ((prevUL * upper propFH-dUL) / (lower CF)) = 1 / ((0.7*0.016) / (1/3,247) ≈ 2.7%) to about 1/7 (calculation upper bound: 1 / ((prevUL * lower propFH-dUL) / (upper CF)) = 1 / ((0.7*0.004) / (1/2,563) ≈ 13.9%) females with an FH deficient UL is expected to carry a germline *FH* variant.

## DISCUSSION

Identifying individuals who carry a pathogenic variant in the *FH* gene is considered important as they have an increased lifetime risk of about 15%(55) for the aggressive renal cell carcinoma (RCC) type associated with HLRCC. A timely diagnosis can enable clinical surveillance for RCC and may benefit these individuals and their families.(55) Due to the distinctive histomorphological appearance of the FH deficiency-driven neoplasms of uterus and kidney, pathologists play a central role in their initial recognition and hence identification of patients with increased risk for such hereditary tumor syndromes. Nevertheless, they are often blinded for the patients’ detailed medical and family history. Cutaneous leiomyomas, especially if multiple, are pathognomonic and FH-deficient RCC in young adults before age 50 years also raises a strong suspicion for HLRCC. Due to the high lifetime risk for ULs in women and the associated frequent need for surgical treatment, the FH deficient UL subtype is frequently encountered in the pathologist routine despite constituting only up to 1.6% of all ULs. However, there is currently no consensus approach to identify the subgroup among patients with ULs who carry a germline *FH*-variant.

Different screening approaches based on morphological or immunohistochemical methods have been recently proposed and tested(15, 17). These studies highlighted the value of routine morphological assessment as strong screening tool assisted by adjunct IHC in the initial recognition of FH-related ULs. Further investigating our previously reported cohort(13), we now characterized the somatic variant status in 13 ULs with distinctive histomorphological features and confirmed FH loss by IHC. By using an established capture-based panel sequencing approach together with comprehensive bioinformatic analyses for SNVs/indels and CNVs, we could identify biallelic variants in all 13 analyzed cases. In eight cases, we identified a SNV/indel together with a CN-loss on the second allele, while the remaining five cases showed biallelic CN-losses. The observation that no UL had two SNVs/indels, points to an initial *FH* mutation (SNV/indel or CNV) in a progenitor cell which then increased the probability for a CN-loss in descendent cells. Fumarase is known to play a role in response to double strand-breaks (DSB) by activating the non-homologous end joining (NHEJ) mechanism(56). When the NHEJ pathway is inactivated the DSB are repaired by more error prone mechanisms, like microhomology-mediated end joining, which can lead to deletions. The proportion of SNVs/indels detected in our study is comparable to previous reports(14, 16, 17, 57). However, the authors of these publications could often not identify CN-loss of the second allele since the CN-analysis and the sequencing technique used, did not allow to reliably estimate the allele frequency. Interestingly, Joseph and colleagues used a similar approach to identify a biallelic CN-loss in one FH deficient UL which had escaped Sanger based mutation detection before(17). The detection/identification of 5/13 ULs with biallelic CN-loss and 8/13 monoallelic CN-loss in our study, suggests CN-loss as a predominant mutational mechanism which has been underestimated, but can be reliably detected using panel sequencing. The combination of CN- and VAF-analyses not only allowed us to detect both *FH* mutations in all 13 tumors, but also raised the suspicion of intratumoral heterogeneity in one case (S05), a mechanism only recently proposed to further complicate detection of the cause of FH-associated ULs(15).

Notably, our series of ULs had been preselected based on complete FH-loss in IHC. While biallelic CN-losses or the combination of a CN-loss of one allele and a truncating variant on the other allele (S03: splice-acceptor variant, S06: frameshifting variant) are expected to result in the lack of protein product, missense variants usually do not cause reduced protein dosage but exchange only single amino acid residues. Rabban and colleagues reported normal FH immunoexpression in tumors of two females with pathogenic germline *FH* missense variants(15). We observed no difference in IHC and morphology for ULs with biallelic deletions, likely gene disrupting and missense variants in *FH.* This is clearly exemplified in Figure 3 for the missense variant c.587A>T, p.(His196Leu), affecting the in our cohort recurrently mutated amino acid residue H196, in comparison to both a case with biallelic deletion and the truncating variant c.457delG, p.(Val153*). The observation of six variants annotated as missense was thus unexpected, also as the majority of (likely) pathogenic mutations (59.3%) in databases are either truncating or affect the splice-sites (Figure S1) and identifying six missense variants out of 8 total variants is unlikely (Binomial p-value 0.045). A possible explanation would be that the six missense variants we identified in fact pose a different effect on the gene product by, for example, altering the mRNA splicing as has been described for other exonic variants(58). The predicted missense variant c.1236G>A, identified in individual S09, affects the last base of exon 8 and can be expected to result in aberrant splicing. For the other five variants though, algorithms indicated normal splicing behavior, leaving another mechanism to explain loss of protein. We therefore utilized a published crystal structure to perform a three-dimensional variant clustering, which showed that the distribution of missense variants in our samples are unlikely by chance (Figure 1B; Figure S2). Hence, it seemed reasonable that the protein domain was important for proper FH expression. A literature search showed that other missense variants (c.922G>A, p.(Ala308Thr) or A308T; c.952C>T, p.(His318Tyr) or H318Y) affecting the same domain result in defective fumarase oligomerization(59) and we could show that these two variants significantly colocalize with the missense variants identified herein (Figure S2B). Interestingly, a case report showed loss of FH IHC staining in an individual with another missense variant (c.953A>T, p.(His318Leu)) affecting the same His318 amino acid residue(60). Altered tetramer formation and a “dominant negative” effect has been shown for one of the most frequently described *FH* “hot spot” mutations (c.698G>A, p.(Arg233His) or R233H and often referred to as R190H)(61) and cases with this variant have been shown to have FH loss in IHC(62). Together our results and the literature indicate altered oligomerization of FH through certain missense variants affecting subunit interactions as a mechanism causing loss of FH expression in IHC(59, 63). A possible explanation for this observation is reduced stability through by degradation of non-tetrameric fumarate hydratase, a hypothesis which should be further investigated. These results are particularly interesting as the combination of classical pathology techniques with molecular genetics and bioinformatic analyses pinpointed the specific function of certain variants in a protein domain. This indicates that larger data collections and systematic evaluation will yield novel hypothesis and insights, which can then be evaluated by functional studies. Nevertheless, our finding of a specific missense cluster causing loss of FH expression also indirectly confirms the concerned observation that other missense variants will be missed by screening methods based on FH IHC. This might be avoided by using 2SC IHC when it is clinically available.

**Figure 3.**
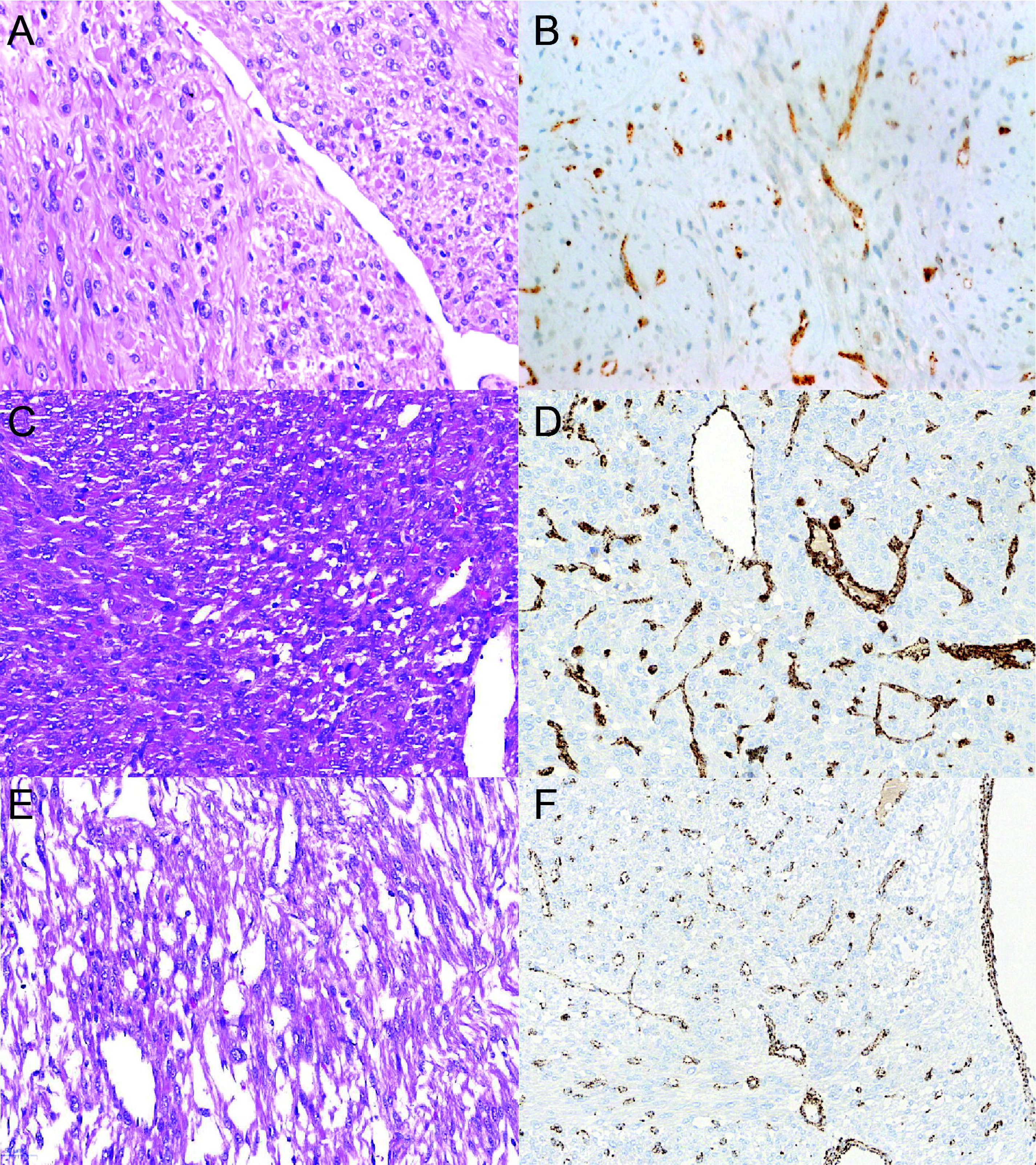
Exemplary histopathology and FH IHC. Exemplary FH IHC staining and morphologic features for ULs with biallelic deletions, likely gene disrupting and missense variants. (A, B) S12 called with both a 15.9 kb and a 1407.7 kb deletion. (C, D) S06 with the likely gene disrupting variant c.457delG, p.(Val153*) and a large 42,557.9 kb deletion. (E, F) S05 with the missense variant c.587A>T, p.(His196Leu) and a small 15.9 kb deletion. Note the similar immunohistochemical FH loss and morphology in all three samples despite the diverse mutation types. Endothelial cells with intact FH expression serve as internal positive control.

Despite the successful complete characterization of the dual *FH*-hits in all 13 ULs, the question remained whether the identified aberrations are somatic, indicating sporadic disease, or in fact are germline and potentially associated with HLRCC. The ethics consent for this study did not allow us to test adjacent normal tissue or to recontact the patients regarding germline testing. Instead, we mathematically estimated that the observed VAF in the eight ULs with an SNV/indel are best explained by somatic two hit model. As we could not directly estimate tumor purity from the sequencing data and VAFs of somatic variants due to the low overall mutational load, we cannot fully exclude that some of the identified eight SNVs/indels and maybe also CNVs in the remaining five cases are indeed germline variants. However, our reasoning that all identified variants are somatic is additionally supported by our calculation that only 1/36 (2.7%) up to 1/7 (13.9%) of females with an FH deficient UL are expected to carry a germline *FH* variant. Thus, it is not unlikely (Binomial p-value = (1-0.139)^13 ≈ 0.143) to find no germline carrier in the 13 cases analyzed in our study. In this regard, the screening strategy proposed by Rabban and colleagues, based on FH deficient UL morphology, is especially interesting as it allowed identification of a germline variant in 5/2,060 individuals(15) which equates to a enrichment of germline carriers between 6.2x (≈ (5/2,060) / (1/2,563)) and 7.9x (≈ (5/2,060) / (1/3,247)). Nevertheless, statistical performance measures of this approach can currently not be estimated reliably.

Panel sequencing further allowed us to identify two somatic likely gene disrupting mutations in *TP53* in sample S04, which is comparable to a homozygous *TP53* deletion described in a UL sample with FH-loss in IHC and a missense mutation with loss-of-heterozygosity(16). While IHC showed a reduced expression of RB1 in all samples analyzed, CN-analysis revealed a likely heterozygous loss of the *RB1* gene locus in 25% of the examined ULs. This finding is comparable to the results of Bennett and colleagues who identified homozygous *RB1*-losses in 40% of ULs with normal FH staining but not in FH deficient ULs(16). Complete RB1-loss is therefore rather a feature of FH-normal ULs. Additionally, IHC showed variable low expression of RB1, hardly distinguishable from complete loss, in all analyzed samples which excluded it as a routine diagnostics marker. Our CN-analysis identified an enrichment of CN-gains at *ALK*-gene locus previously not reported in ULs. This observation is interesting as it could offer novel treatment options with *ALK*-inhibitors(64) but needs independent confirmation. While our analysis of mitochondrial genome dosage and telomere content from panel sequencing data did not identify significant differences compared to other tumors (Supplementary notes; Figure S6), it confirms the added value of next-generation based sequencing to investigate novel hypotheses from available data.

Finally, further systematic sequencing analyses like the initial studies of Mehine and colleagues(11, 65) on larger cohorts of ULs will be required to fully define the mechanism involved in the development of these benign uterine tumors causing significant morbidity in the female population and to investigate associations with heritable tumor syndromes like the HLRCC. We anticipate that this will first require unbiased tumor/normal sequencing (panels, exomes or genomes) but also RNAseq and methylation studies in a representative cohort. Future interesting fields of investigation would be individuals with multiple ULs who do not have a germline variant, where sequencing of multiple tumors from the same individual might uncover somatic mosaicism.

In conclusion, the combination of IHC screening and panel sequencing with detailed bioinformatic analyses allowed the identification of both genetic hits in all the 13 ULs studied, confirming the established Knudson hypothesis in *FH*-related tumor development and the role of *FH* as a tumor suppressor gene. This successful approach allowed us to identify a cluster of missense variants associated with immunohistochemical protein reduction, proving that missense variants contribute to FH deficient ULs. We agree with previous concerns(15, 17, 18) that some pathogenic missense variants in individuals with HLRCC might be missed using only FH IHC as screening method. The proposedly more reliable 2SC IHC is currently not available for routine use. While screening of certain morphologic features in tumors has been shown to enrich for patients with HLRCC(15), the statistical performance of this approach can currently not be assessed reliably. Thus, a prospective tumor/normal sequencing study, which represent the current gold-standard(17), in a representative risk group is needed. Based on literature recommendations(55, 66) and our experience with heritable tumor syndromes, a possible strategy would be to offer this tumor/normal screening to every symptomatic woman below age of 40 years who has multiple ULs or one UL ≥ 10cm in diameter without prior selection based on morphological features. Further inclusion criteria could be a positive personal or family history for tumors, if this information is available to the pathologist. The use of a relatively small but standardized commercial gene panel for screening, like in this study, can reduce the associated costs and allow inter-institutional data collection and collaboration between pathologists and geneticists.

## SUPPLEMENTARY

**Supplementary notes |** Eight additional figures, three additional tables and supplementary methods detailing the results and providing further information about the analyses performed. Additionally, the case report of individual S11 is described and discussed in detail together with methods used to confirm pathogenicity of the identified germline *TSC1* variant.

**Supplementary File S1 |** Excel file containing the worksheets “summary”, “cases”, “panel_stats”, “trusight_cancer_v1_genes”, “FH_SNVindel_summary”, “SNVindel_all”, “FH_CNVkit_summary”, “CNVkit_aberrations”, “mitochondria” and “telomere”. The “summary” worksheet contains a detailed description of all other worksheets and the respective data columns.

**Supplementary File S2 |** Excel file containing the worksheets “summary”, “FH_domains”, “allFH_missense”, “FH_ClinVar”, “FH_LOVD”, “FH_COSMIC”, “FH_gnomAD”, and “FH_BRAVO”. The “summary” worksheet contains a detailed description of all other worksheets and the respective data columns.

## ACKNOWLEDGEMENTS

The authors would like to thank Rudolph Jung, Christa Winkelmann, Andrea Eberwein and Leonora Klassen for excellent technical assistance. B.P. is supported by the Deutsche Forschungsgemeinschaft (DFG) grant PO2366/2-1.

## COMPLIANCE WITH ETHICAL STANDARDS

The authors confirm that the study has been performed in accordance with accepted principles of ethical and professional conduct for biomedical scientific research. This study is covered by the ethical vote for retrospective translational research studies of the Ethical Committee of the Medical Faculty of the Friedrich-Alexander-Universität Erlangen-Nürnberg.

## SUPPORTING INFORMATION

Supplementary information is available at the publisher’s website.

