## Supplementary notes for "Targeted sequencing of FH-deficient uterine leiomyomas reveals biallelic inactivating somatic fumarase variants and allows characterization of missense variants"

Running title: **Biallelic Mutations in FH deficient ULs**

^1^Institute of Human Genetics, University Hospital Erlangen, Friedrich-Alexander-Universität Erlangen-Nürnberg (FAU), Erlangen, Germany

^2^Institute of Human Genetics, University of Leipzig Hospitals and Clinics, Leipzig, Germany

^3^Institute of Pathology, University Hospital Erlangen, Friedrich-Alexander-Universität Erlangen-Nürnberg (FAU), Erlangen, Germany

^4^Department of Obstetrics and Gynecology, University Hospital Erlangen, Comprehensive Cancer Center ER-EMN, Friedrich-Alexander University Erlangen-Nürnberg (FAU), Erlangen, Germany

^*, #^These authors contributed equally to this work

Correspondence to:

André Reis, Institute of Human Genetics, University Hospital Erlangen, Friedrich-Alexander-Universität Erlangen-Nürnberg (FAU), Schwabachanlage 10, 91054 Erlangen, Germany,, phone: +49 9131 8522319, fax: +49 9131 23232

Abbas Agaimy, Institute of Pathology, University Hospital Erlangen, Friedrich-Alexander-Universität Erlangen-Nürnberg (FAU), Krankenhausstrasse 8-10, 91054 Erlangen, Germany,, phone: +49 9131 8522288, fax: +49 9131 8524745

**SUPPLEMENTARY MATERIALS AND METHODS**

**Mitochondrial genome dose calculation**

Mitochondrial gene dosage in the 13 ULs was calculated as described previously(1) and compared to two in-house control sets (15 HBOC-associated tumors, HBOC; 100 germline controls, CNT) sequenced using the same platform. In brief, the relative mitochondrial genome dosage (MGD; ratio chrM/chr2) was calculated by dividing the coverage on the chrM contig of the hg19 human reference by the on-target coverage of chromosome 2 (chr2 in hg19, used because of the deletions affecting the *FH* gene on chr1). Coverage was calculated using Qualimap2(2) using the BED-file for the TruSight Cancer Sequencing Panel v1 kit(3) provided by Illumina (San Diego, USA) with chrM (hg19: chrM:1-16571) added.

**Telomere content calculation**

Telomere content was calculated as described previously(1) using TelomereHunter(4) and compared to the same in-house control sets used for MGD calculation.

**Genetic counseling and germline molecular genetic testing**

After the identification of FH deficiency in one resected UL, genetic counseling and molecular genetic testing for HLRCC at our Center for Rare Diseases was recommended in the pathology report. Genetic consultation included detailed medical, family history and clinical examination by a trained geneticist. Upon informed written consent, molecular genetic testing on DNA derived from peripheral blood lymphocytes was performed by PCR and Sanger sequencing of the *FH* gene with custom primers (Supplementary notes and Table S1) together with multiplex ligation probe amplification (MLPA; Kit P198, MRC-Holland). In individual S11 tuberous sclerosis was suspected clinically and targeted sequencing using the TruSight Cancer Sequencing Panel v1 (Illumina, Inc., San Diego, USA) was additionally performed as described previously(5).

**Sanger sequencing of *FH***

Germline *FH* variant analysis and confirmation of variants identified by panel sequencing in the 13 ULs was performed by PCR and Sanger sequencing using standard procedures and provided in Table S1.

**RNA isolation from blood lymphocytes of S11**

Total RNA was extracted from blood lymphocytes with the PAXgene Blood System (Becton Dickinson, Franklin Lakes, NJ). cDNA was produced with Superscript II Reverse Transcriptase Kit (Invitrogen, Carlsbad, CA) according to the manufacturer's instructions.

**RT-PCR analysis of *TSC1* c.2040A>G**

RNA from PAXgene stabilized fresh blood was used for RT‐PCR and Sanger sequencing of *TSC1* was performed with exon spanning primers (5′‐TCCACATCTGGATGTCCTTCTCTTG‐3′ and 5′‐GAGCAGACGCGCACAGCAAG‐3′) following standard procedures.

**Targeted cDNA sequencing of RNA from S11**

After rRNA depletion and DNA digestion we performed first- and second-strand cDNA synthesis followed by targeted enrichment of cDNA using the TruSight Cancer Sequencing Panel and sequencing on an Illumina MiSeq platform (both: Illumina, Inc., San Diego, USA). Resulting read files were aligned to the hg19 reference genome from the GATK(6) (Genome Analysis Toolkit) bundle and the GENCODE(7) splice junction annotation release v19 using the splice aware read aligner STAR(8) version 2.6.1a in two step-mode. The alignment step was repeated with an alternative hg19 reference with germline variants of S11 from the DNA panel sequencing introduced using ´FastaAlternateReferenceMaker´ from within GATK. GNU Parallel(9) was used for job parallelization and software was installed through Conda (Anaconda Inc., Austin, USA). Resulting alignment files were sorted and indexed with Picard tools(10). BAM files were visualized and inspected for alternative splicing using the IGV(11) browser and its inbuild Sashimi-plot functionality.

**Data analysis and plotting**

The data provided in Supplementary files 1 and 2 (Excel; Microsoft Corporation, Redmond, USA) was analyzed and plotted using R language version 3.5.2 with RStudio IDE version 1.1.463 (RStudio Inc., Boston, USA). Libraries used were "plyr", "tidyverse", "readr", "readxl", "Rmisc", "ggsignif", "ggrepel" and "cowplot". Illustrator and Photoshop CC 2018 (Adobe Systems, San José, USA) or Inkscape 0.92.3 (https://inkscape.org/) were used to adjust main Figure 1 for parts which could not be directly composed in R and to compose main Figure 2 and 3 from primary image data.

**SUPPLEMENTARY RESULTS**

**Mitochondrial genome and telomere content**

Both the *FH* and *TERT* genes encode for moonlighting proteins (fumarase and telomerase) performing functions in the mitochondria and the cell nucleus and associated with cellular aging and tumorigenesis(12, 13). Therefore, we also investigated the telomere content (TC) and mitochondrial gene dosage (MGD) in ULs. While the 13 ULs showed significantly higher TC and MGD, when compared to germline controls (from peripheral blood lymphocytes), the difference was not significant for both analyses when compared with 15 in-house hereditary breast and ovarian cancer (HBOC)-associated tumors sequenced on the same platform (Figure S6).

**Tuberous sclerosis in individual** **S11**

At age 34 years, individual S11 had a symptomatic fibroid of the uterus (causing severe dysmenorrhea) which increased in size for about seven years. The posterior wall myoma was surgically removed followed by reconstruction of the uterus. Histopathological examination revealed atypical leiomyoma and IHC showed deficiency of FH expression. Perivascular epitheloid cell tumor (PEComa) was excluded by presence of histological features characteristic of FH deficient leiomyoma and by lack of melanocytic marker expression. An examination of the kidneys by means of ultrasound as well as the other gynecological examination was inconspicuous. A specialist dermatological investigation was not done, but melanocytic skin nevi were removed by the general practitioner and the histopathological examination of these was unremarkable. At the age of 18 years, she had a seizure disorder which had been treated with lamotrigine and resulted in a seizure free interval for three years. A brain MRI conducted because of dizziness at age 33 years raised the suspicion of multiple sclerosis-like unclassified lesions. Examination of the skin revealed multiple small papules on the forehead, and a larger (2 x 2.5mm) similar change on the scalp. She had one large 8cm hypomelanotic spot on the left lower back and a smaller one on the right upper arm. Iris abnormalities or signs of nail fibromas were not noted. Family history showed that her mother had a double-walled uterus, her maternal aunt had only one kidney, her maternal grandmother had colorectal cancer at age of 88 years and this grandmother’s mother died of renal cancer at about 60 years of age. Her 27-year-old half-brother also had epilepsy. Clinical examination and family history thus raised the suspicion of tuberous sclerosis complex.

Molecular genetic testing by means of Sanger sequencing and MLPA identified no pathogenic germline variant in the *FH* gene. Also, panel-based sequencing of other genes associated with increased renal cancer risk (*SDHB*, *SDHC*, *SDHD*, *SDHAF2*, *PTEN*) was unremarkable. Extended panel analysis identified the synonymous variant c.2040A>G affecting the penultimate position of exon 16 of *TSC1*. Subsequent RT-PCR and Sanger sequencing showed aberrant RNA splicing and confirmed the suspected diagnosis. Tumor panel sequencing in this case showed biallelic loss of *FH* without evidence for a second-hit in the *TSC1* gene.

**SUPPLEMENTARY DISCUSSION**

**Germline *TSC1* mutation and FH deficient UL**

Detection of a *TSC1* germline mutation in an individual (S11) with FH deficient UL in this study is novel. This observation on the one hand highlights the diagnostic strength of genetic counseling combined with broad panel testing in young tumor patients, enabling the identification of rare monogenic tumor syndromes. On the other hand, the association of FH deficient ULs and this *TSC1* variant is intriguing as both fumarase and the tuberous sclerosis proteins hamartin (*TSC1*) and tuberin (*TSC2*) influence HIF1α signaling(14) and heterozygous *TSC1* knock-out mice develop uterine leiomyoma and leiomyosarcoma with subsequent loss of the second allele(15). Perivascular epithelioid cell tumors (PEComas) represent the main uterine manifestation the tuberous sclerosis complex (TSC). These neoplasms overlap strongly with uterine smooth muscle tumors but can be separated by their additional distinctive morphological and immunophenotypic features. In this individual S11, the histology of the tumor (with characteristic features of FH deficient UL; Figure S8) and lack of melanocytic marker expression precluded a diagnosis of PEComa and confirmed instead UL with FH deficiency. Also, while we could confirm aberrant splicing of the identified *TSC1* germline variant and thus its pathogenicity, the VAF in the sequenced UL from the individual showed no significant deviation from 50% and did therefore not further confirm a causal relationship. Currently, no data is available about FH status in uterine PEComa. Notably, a recent report described renal PEComa in a patient with proven HLRCC syndrome and the PEComa was initially judged as a potential RCC on screening of the kidney(16). FH expression was retained in that PEComa indicating an alternative pathogenesis. However, taken together, the current and that previous report suggest a possibility of interaction between the *FH* and *TSC1* gene mutations in rare cases and it remains to be clarified if these two genes might on occasion represent alternate mechanisms in the pathogenesis of rare smooth muscle neoplasms.

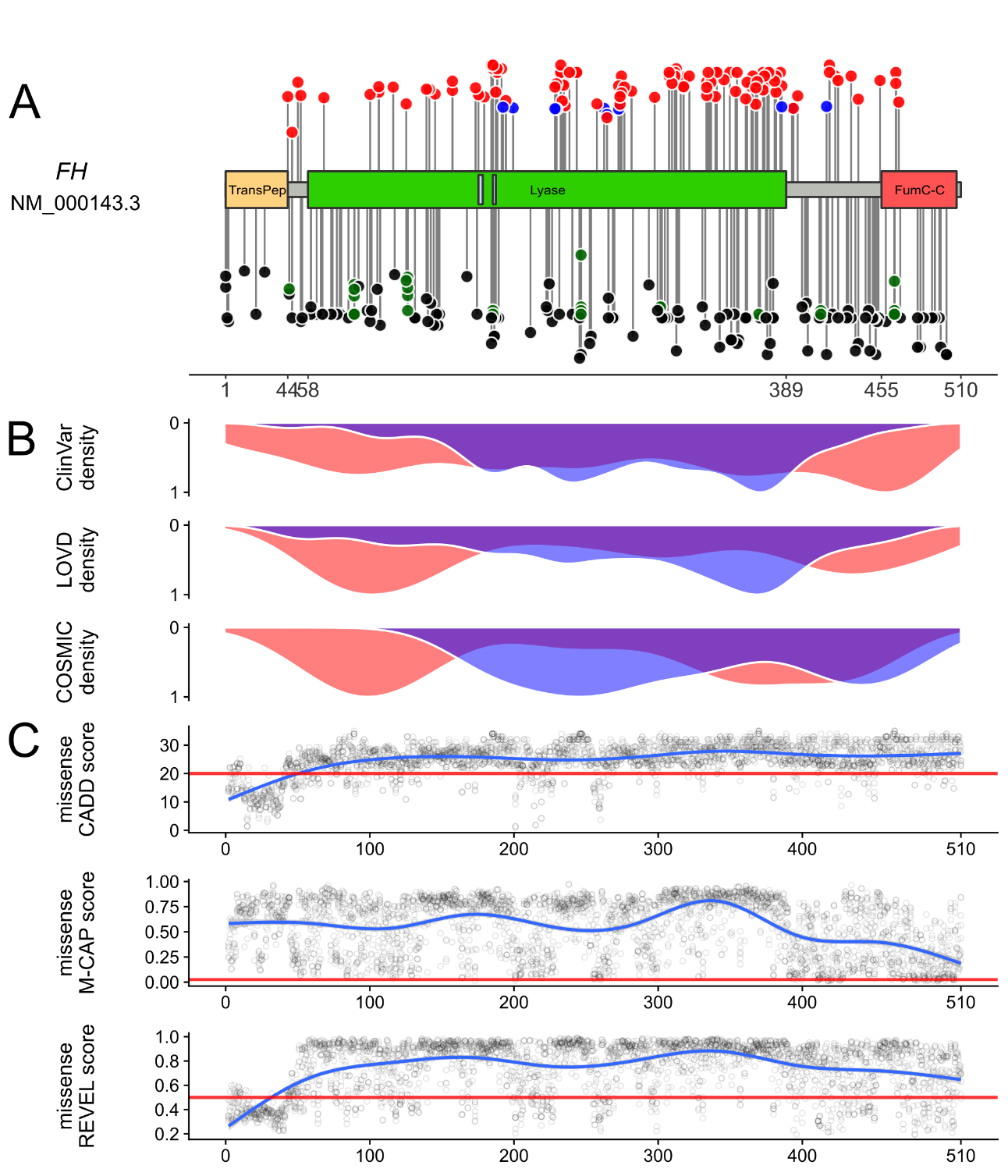
**SUPPLEMENTARY FIGURES AND TABLES**

**Figure S1 | *FH* variant spectrum in databases and domain structure**

**(A)** Schematic representation of the FH protein, domains and localization of (likely) pathogenic variants from the ClinVar(17), LOVD(18, 19) and COSMIC(20, 21) databases. Loss-of-Function (LOF) variants are presented in black and splice-site variants in dark green (below the protein schematic), while missense variants are in red and in-frame insertions/deletions in blue (above the protein schematic). **(B)** Density plot of (likely) pathogenic truncating (red) and missense (blue) variants reported in ClinVar, LOVD and COSMIC databases. **(C)** Generalized linear models (GLM) of the CADD(22), M-CAP(23) and REVEL(24) computational prediction scores for all possible *FH* missense variants. It is evident that missense variants are found mainly in the Lyase domain with a dense cluster in the C-terminal part of this domain. This region correlates with a peak in the GLM curve for all three missense prediction scores, indicating a critical functional region in the protein which is sensitive to missense variation. Note that likely gene disrupting (LGD) variants in contrast are scattered throughout the protein (the peak in the N-terminal region is likely caused by the enrichment of splice variants in this region). See also main Figure 1 and Supplementary data file 2 sheets “FH_ClinVar”, “FH_LOVD”, “FH_COSMIC” and “allFH_missense”.

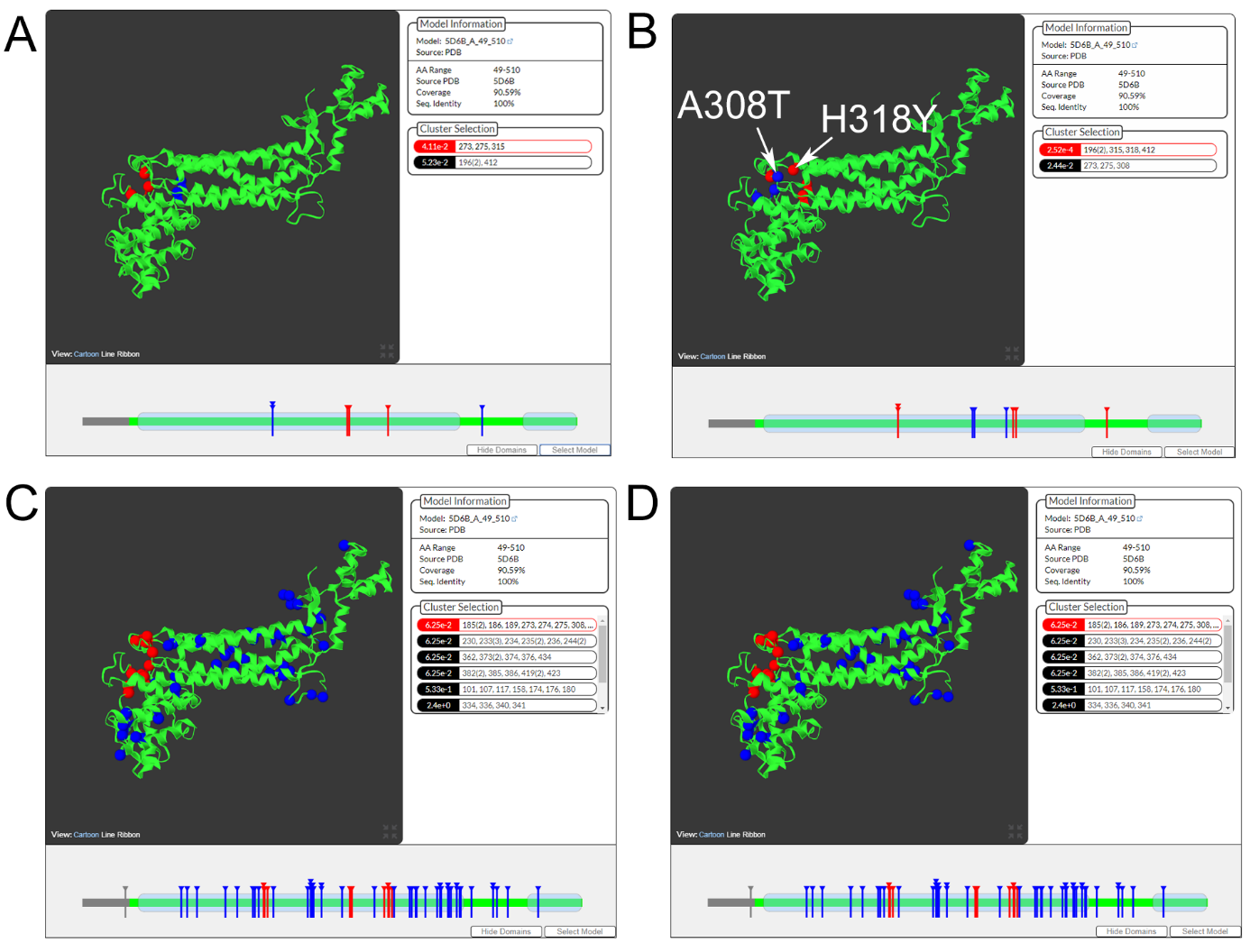

**Figure S2 | 3D clustering analysis**

Screenshots from the 3D clustering analyses using mutation3D(25). Spheres mark amino acid positions of the respective missense variant in the FH protein. Red spheres mark the cluster with the highest p-value in the analysis. **(A)** The 6 herein identified *FH* missense variants (L315P, H196P, G275A, M412I, A273T, H196L)(26), **(B)** the 6 herein described missense variants with the two missense variants A308T and H318Y (marked by white arrows) known to cause defective oligomerization(27), **(C)** the 58 (likely) pathogenic *FH* missense variants(28) reported in ClinVar and **(D)** the 71 (likely) pathogenic *FH* missense variants(29) reported in LOVD. Similar to main Figure 1 the tertiary protein structure data of human fumarase with the PDB-ID: 5D6B(30) was used as template. The missense variants L315P, G275A, A273T identified in sample S01, S07 and S10, respectively, form a cluster with significant p-value (0.0411, empirically derived by mutation3D using a bootstrapping approach). Interestingly, this protein region always forms a cluster (red spheres in A, B, C) with the highest significance p-value in both the ClinVar (B, p-value = 0.0625) and LOVD (C, p-value = 0.0625) dataset, indicating a mutational hot-spot and a specific disrupting effect of variants affecting this protein region. Also note that when adding the two A308T and H318Y causing defective FH oligomerization(27) to the 6 variants annotated as missense from our 13 ULs, these 8 variants form two neighboring clusters with significant p-values (0.000252 and 0.0244) indicating a specific function of this protein domain in oligomerization.

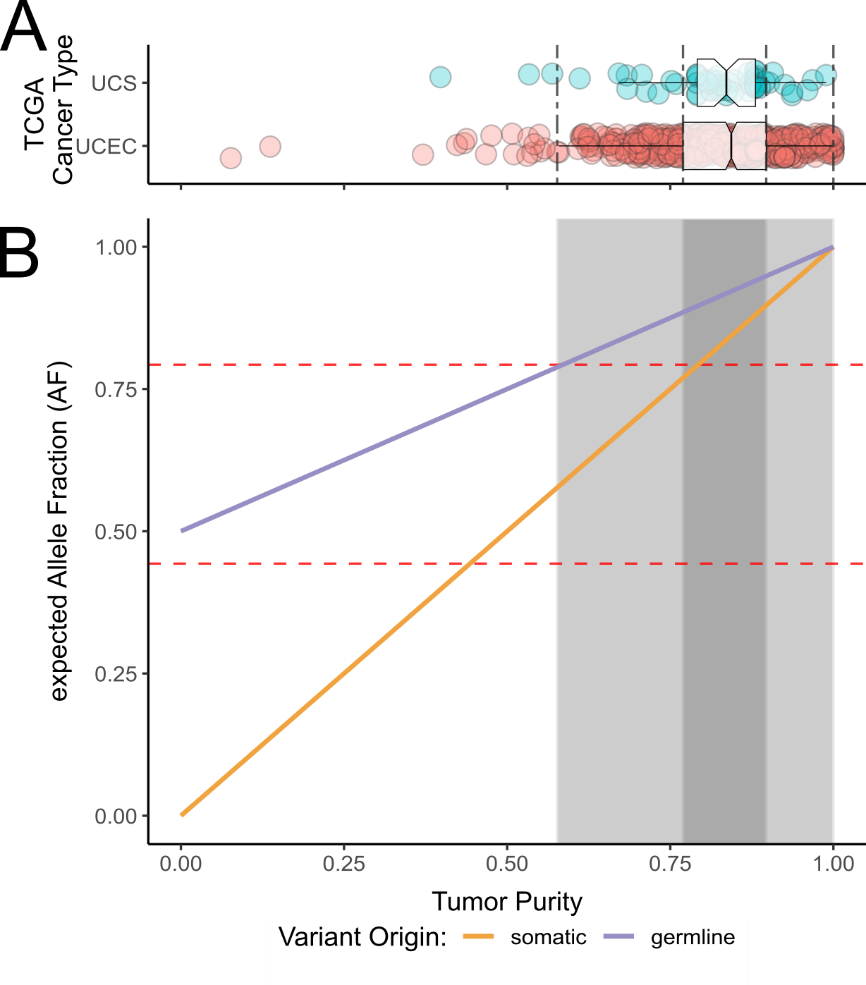

**Figure S3 | Purity models used to assess possible germline or somatic variant status**

**(A)** Box- and scatter-plots of the purity estimates of two uterine tumor types from the TCGA(31) (blue = UCS: uterine carcinosarcoma; red = UCEC: uterine corpus endometrial carcinoma(32)) as published by Aran et al.(33). The 1st (0.770) and 3rd (0.897) quartiles and the boundaries of the whiskers (coefficient = 1.58; lower: 0.577; upper: 1.000) for UCEC are marked by vertical dashed lines to indicate typical purity spreads. **(B)** Line-plot showing the theoretical relationship between tumor purity (TP; defined as fraction of tumor cells) and expected allele fraction (AF) if the observed variant was initially germline or somatic and a somatic second-hit deleted the second allele based on the formula “AF = TP * k + (1 - TP) * l” (k = correction coefficient for non-diploid loci, set to 1 in this simple model; l = initial zygosity in the germline, set to 0.0 for somatic [orange line] or 0.5 for germline [blue line] variants, respectively). The grey areas mark the purity spreads estimated in (A) and the red dashed horizontal lines mark the lower (S05: AF = 0.443) and upper (S06: AF = 0.793) boundaries of the AF observed in the 13 ULs. Note the orange line crossing the grey areas for a large range of AF values, indicating that somatic variants can likely explain the AFs observed in the 13 ULs. Germline variants would be expected to produce higher AFs in the tumor when assuming loss of the second allele.

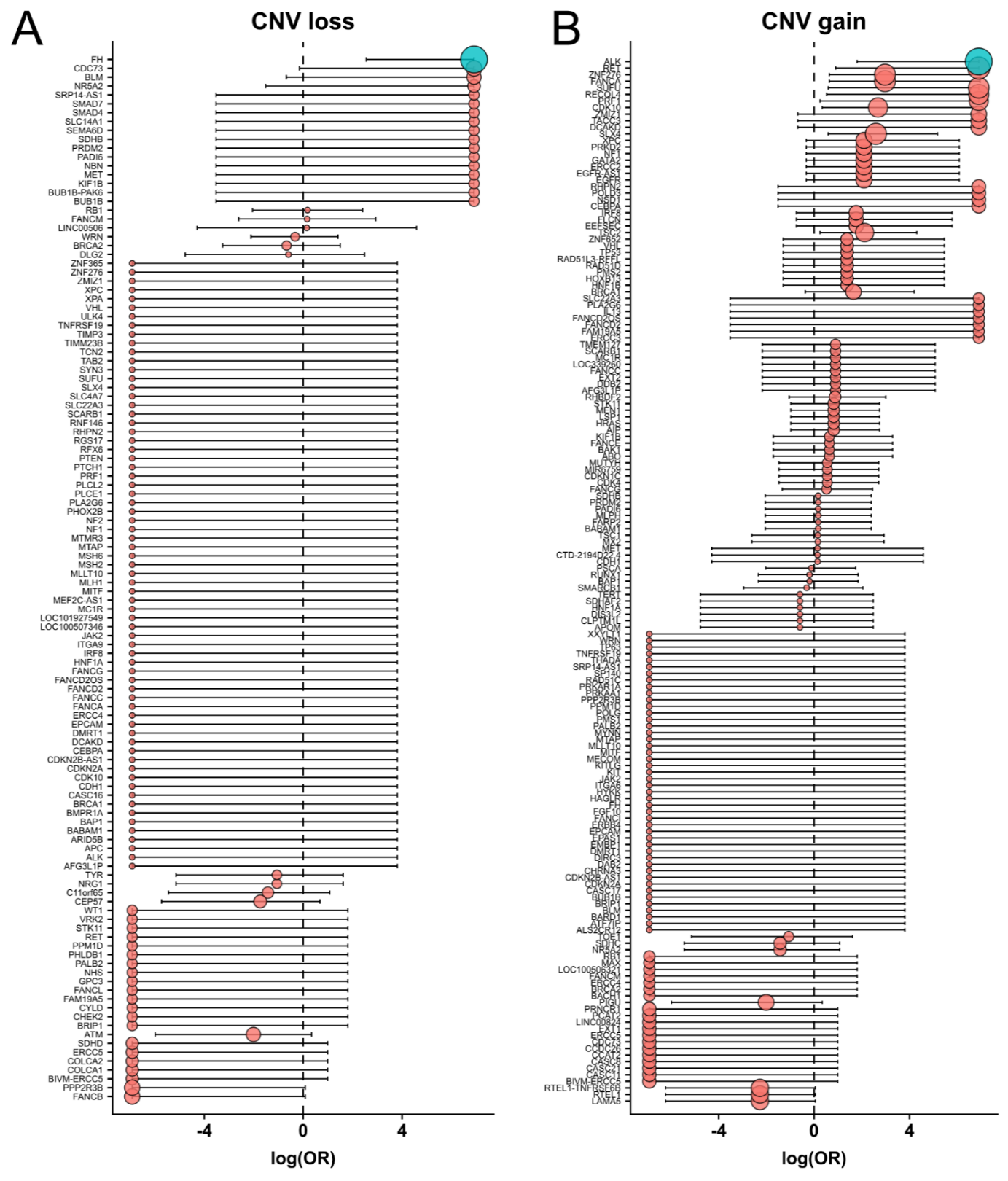
**Figure S4 | CNV enrichment comparing *FH*-deficiency ULs and HBOC controls**

To analyze UL-specific copy number (CN) alterations, we compared the CN-calls of the 13 ULs to 15 in-house HBOC-associated (hereditary breast-ovarian cancer) tumors sequenced on the same time with the same platform and also called with CNVkit(34) using the same settings. For each gene we counted how many samples had a CN-loss (CN < 2) or -gain (CN > 2) sized ≥ 500kb and calculated 2-sided Fisher's exact test statistics. Odds-ratios (OR) were capped at 0.001 and 1000 (log_e_(0.001) ~ -6.90, log_e_(1000) ~ 6.90) and p-values were PHRED-scaled (-log_10_(p)) for visualization. We applied Bonferroni correction for p-values using the total count of genes tested (loss: 0.05 / 118 ~ 0.000424; gain: 0.05 / 159 ~ 0.000314). Genes are sorted by p-values and confidence-intervals, colored dots mark the respective OR and the dot-size is proportional to the PHRED scaled p-value with blue dots indicating significant values while red dots are not significant after correction for multiple testing. Only the *FH*-gene is significant for CN-losses **(A)** and the *ALK*-gene for CN-gains **(B)**. See also Supplement data file 1 sheet “CNVkit_aberrations” for detailed CN calls of both UL and HBOC groups.

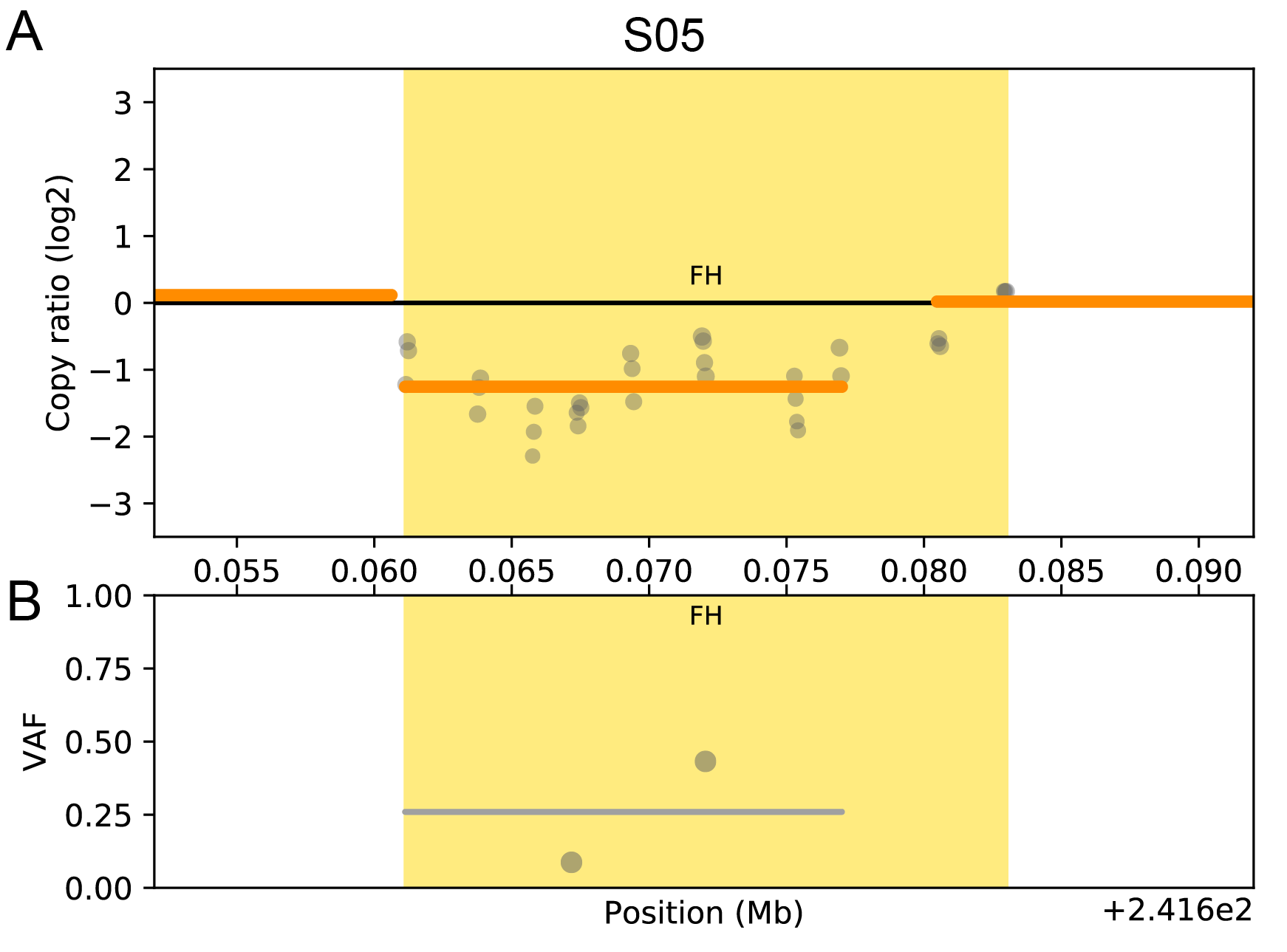

**Figure S5 | CR-profile of sample S05 at the FH-gene locus**

**(A)** CR profile for sample S05 at the *FH*-gene locus with **(B)** VAF (variant allele frequency) plot showing the small 15.9 kb deletion affecting exons 3 to 10 of the one allele.

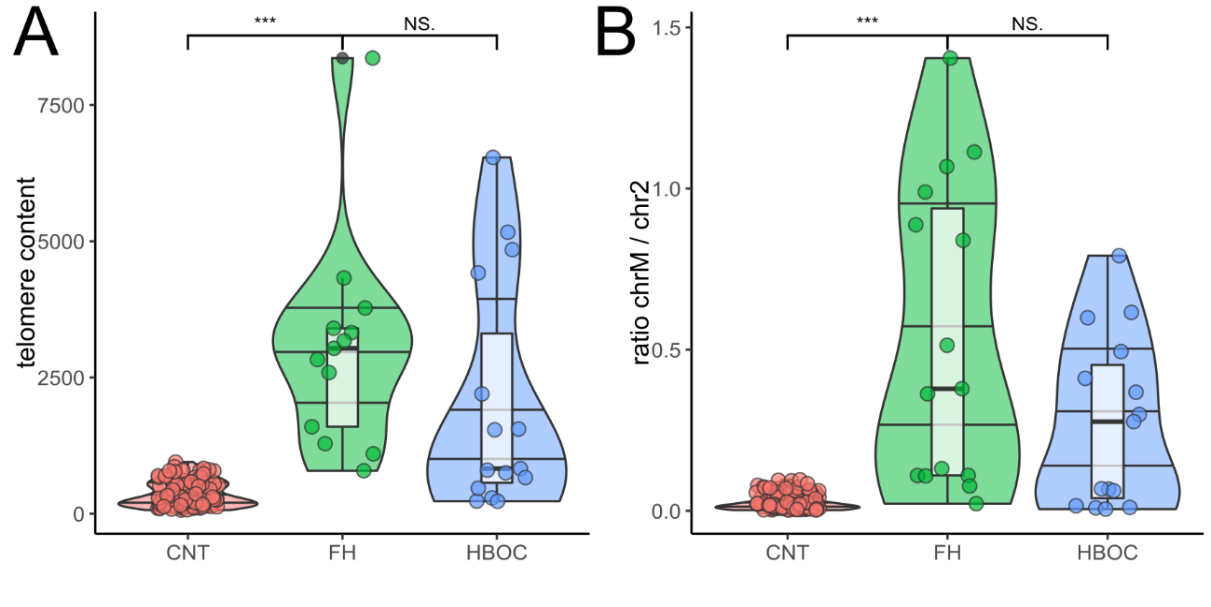
**Figure S6 | Mitochondrial genome and telomere content comparison**

**(A)** Telomere content (TC) was computed from panel-data using TelomereHunter(4). While the 13 ULs show significantly higher TC when compared to germline CNT, the difference between ULs and HBOC controls is not significant. **(B)** The 13 ULs show significantly higher relative mitochondrial genome dosage when compared to germline CNT, but the difference between ULs and HBOC controls is not significant (2-sided Wilcoxon signed-rank test used for statistical comparison). Data from Supplement data file 01 sheets “telomere” and “mitochondria” plotted as scatter and violin plots with box-plots.

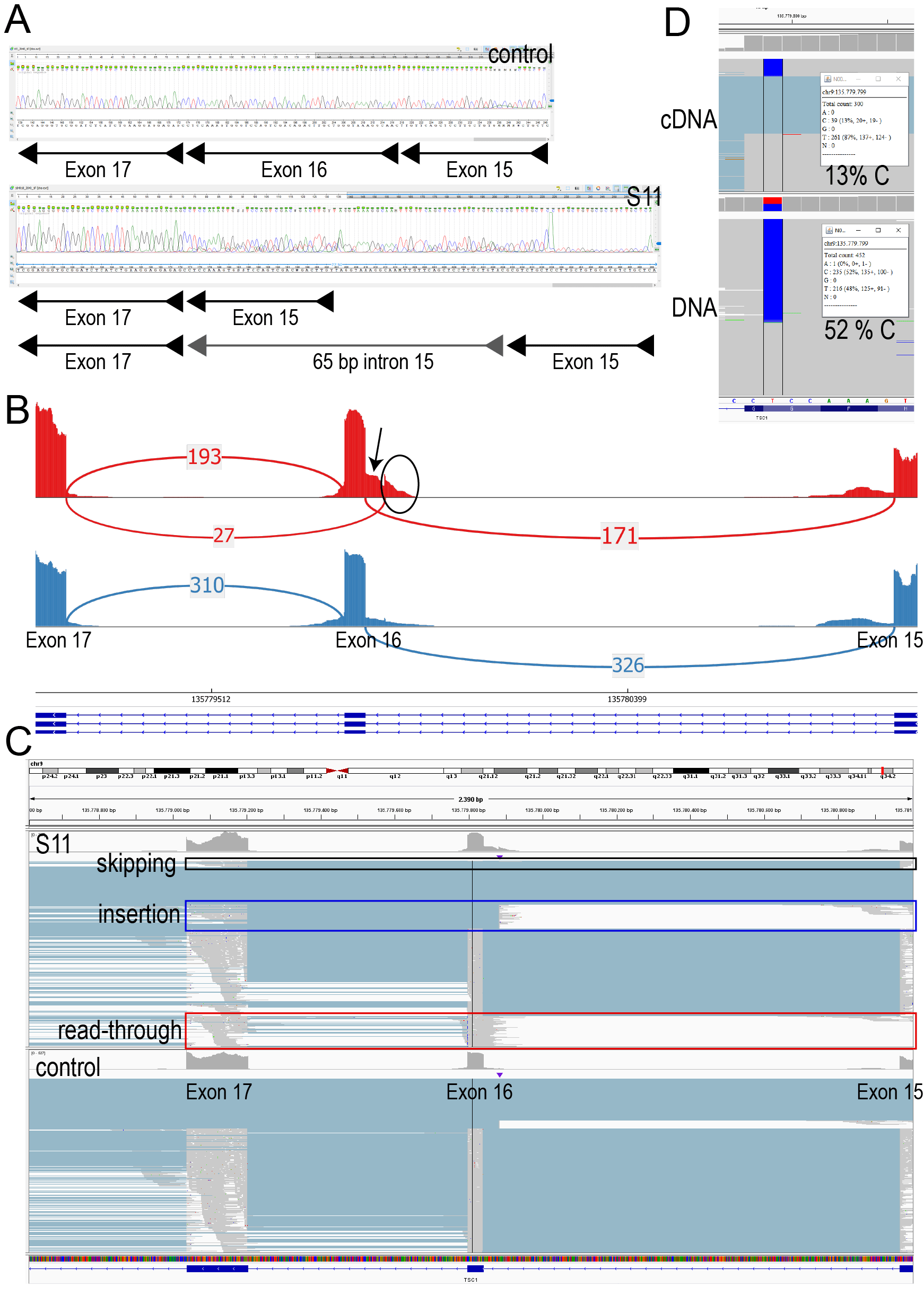

**Figure S7 | Splice effect of the *TSC1* germline variant c.2040A>G in S11**

**(A)** RT-PCR and subsequent Sanger-sequencing (Primers 5′‐TCCACATCTGGATGTCCTTCTCTTG‐3′ and 5′‐GAGCAGACGCGCACAGCAAG-3′) on cDNA derived from PaxGene RNA of individual S11 and wildtype control. The control shows only the expected wildtype splice-product (exon 15 - exon16 - exon 17) while in S11 two other products are present. By subtracting the ambiguity code and comparing with the reference we could determine the smaller product to result from skipping of exon 16 (r.1997_2041del) while the larger product resulted from skipping of exon 16 with insertion of a 65 bp fragment from intron 15 due to the activation of cryptic splice-sites (r.[1997_2041del; 1997_2041ins1997+1021_2041-42]). Both these aberrant mRNA transcripts are expected to result in truncated TSC1 proteins (p.Pro668Serfs*5 and p.Leu667_Ser1164delinsPhe). **(B)** Sashimi-plot of targeted cDNA sequencing (red = S11, blue = control) confirms the RT-PCR results and shows splicing to the cryptic exon (e.g. absent in the normal but included in the aberrant transcript) in intron 15 with skipping of exon 16 (marked by ellipse). Additionally, there is evidence of an intronic read-through transcript (marked by arrow), previously missed by RT-PCR due to the likely very large size of this aberrant transcript which cannot be amplified by standard RT-PCR. Note that exon 16 skipping is not evident in this Sashimi-plot, because the aberrant junction was detected in less 10% of the reads covering the adjacent exons and thus filtered for visualization (see also Figure S7C). **(C)** Visualization of the different aberrant transcripts in IGV browser (upper panel S11, lower panel = control) showing evidence for the exon 16 skipping transcript (black box), the exon 16 skipping and intron 15 insertion transcript (blue box) and the intronic read-through transcript (red box). Note the relatively low abundance of all aberrant transcripts in (B) and (C) indicating nonsense-mediated mRNA decay of these transcripts. **(D)** Comparison of variant allele fraction (VAF) at the *TSC1* c.2040A>G genomic position (chr9(hg19):g.135779799T>C) in the cDNA (13% C-bases) and DNA (52% C-bases) panel of S11 further confirms decay of the aberrant transcripts and preferential expression of the wildtype allele. Note that 13% VAF in the cDNA sample is likely an overestimate as these are mostly the read-through transcript and residual DNA.

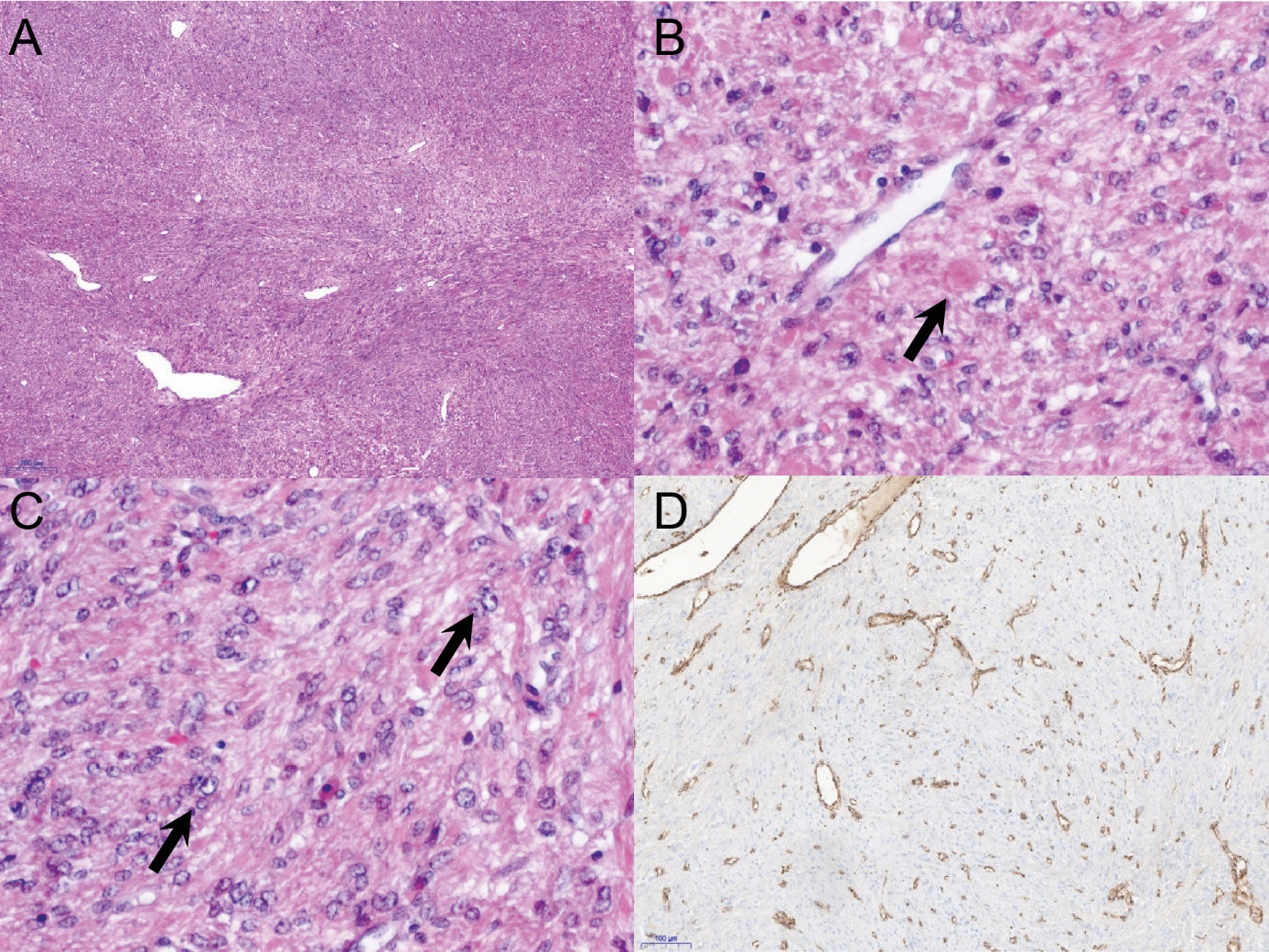

**Figure S8 | Histopathology and IHC of case S11**

Exemplary morphologic features and IHC staining of an UL from individual S11 with the germline *TSC1* variant c.2040A>G. (A)-(C) S11: Histomorphology of FH deficient UL, H&E (note the hyaline globular bodies in (B; arrow) and perinucleolar halos in (C; arrow); (D) S11 with immunohistochemical FH loss (FH IHC).

| NAME(35) | *FH* Exon | LEFT_PRIMER | RIGHT_PRIMER | Chr | Start | End | PRODUCT SIZE | UCSC In-Silico PCR | MeltingTemp F | MeltingTemp R | dbSNP138_F | dbSNP138_R |
| --- | --- | --- | --- | --- | --- | --- | --- | --- | --- | --- | --- | --- |
| FH_e01p | Exon01 | TGTGAGGCTGTTGATTGGAT | GGAGGGCTGAAGGTCACTG | chr1 | 241682839 | 241683137 | 299 | ok | 59.1 | 60.8 | ok | ok |
| FH_e02p | Exon02 | AAGATGCGATTACTTTTGATCC | TGAATACAGCCTACTTCATCCAA | chr1 | 241680428 | 241680683 | 256 | ok | 57.4 | 58.8 | ok | ok |
| FH_e03p | Exon03 | CTGCCAAAATAATAAACTTCCATGC | GCCAGAGCATATCGTCATCC | chr1 | 241676830 | 241677083 | 254 | ok | 62.0 | 60.6 | ok | ok |
| FH_e04p | Exon04 | AAACTCTGTGGCATAATCAGCA | CAAGAACAATCTCAGGTATGCTTT | chr1 | 241675203 | 241675511 | 309 | ok | 59.8 | 58.9 | ok | ok |
| FH_e05p | Exon05 | GCTGGGTTTTGAGTAGTTAGTTGG | GGCCATTTGTACCAAGCTCT | chr1 | 241671852 | 241672200 | 349 | ok | 60.4 | 59.2 | ok | ok |
| FH_e06 | Exon06 | CCTCATCCTTCCCTATACTTTGC | CAAGAATTCAAGACAGGAACACTC | chr1 | 241669215 | 241669647 | 433 | ok | 60.3 | 59.3 | ok | ok |
| FH_e07 | Exon07 | ATTGGAACTTTCTGTTTCACTTG | TTTAAACAAAGAGAGACATGGTCC | chr1 | 241667254 | 241667632 | 379 | ok | 57.5 | 58.7 | ok | ok |
| FH_e08 | Exon08 | TGGGCCTTGCTTTATTGTATATC | TTATGTCACCCAACTACCCAATG | chr1 | 241665633 | 241665983 | 351 | ok | 59.4 | 60.9 | ok | ok |
| FH_e09p | Exon09 | CATGTTGCCTTAGTAATGTCTCTCTC | TGCTGTTCTCAAACACTGATCC | chr1 | 241663642 | 241663958 | 317 | ok | 59.8 | 60.3 | ok | ok |
| FH_e10p | Exon10 | CAATTATGTCACCTTTTGCTTTAGG | GCAGTTTCCTTTCAAACTTATCC | chr1 | 241661035 | 241661346 | 312 | ok | 60.3 | 58.4 | ok | ok |

**Table S1 | *FH*-gene primer pairs used for diagnostics and variant validation**

Primer pairs are based on Kiuru et al.(35) and have been tested by UCSC in-silico PCR, to not lie on a common polymorphism from dbSNP version 138 and validated for diagnostic PCR and Sanger sequencing.

| ID | HGVS_C | HGVS_P | AC | AC norm | Allele number | Reviewed ACMGclass |
| --- | --- | --- | --- | --- | --- | --- |
| gnomad0671 | c.40dupC | p.Leu14fs | 4 | 7,77 | 145674 | Likely pathogenic |
| gnomad0690 | c.2T>G | p.Met1? | 3 | 5,82 | 145704 | Pathogenic |
| gnomad0686 | c.7C>T | p.Arg3* | 1 | 1,93 | 146724 | Likely pathogenic |
| gnomad0692 | c.1A>G | p.Met1? | 2 | 3,18 | 177978 | Likely pathogenic |
| gnomad0408 | c.556-2A>T | NA | 1 | 1,13 | 249830 | Pathogenic |
| gnomad0033 | c.1475_1476delTC | p.Leu492fs | 1 | 1,13 | 250528 | Pathogenic |
| gnomad0407 | c.557G>A | p.Ser186Asn | 1 | 1,13 | 250536 | Likely pathogenic |
| gnomad0406 | c.560C>G | p.Ser187* | 1 | 1,13 | 250648 | Pathogenic |
| gnomad0263 | c.923C>G | p.Ala308Gly | 1 | 1,13 | 251026 | Likely pathogenic |
| gnomad0222 | c.1083_1086delTGAA | p.Glu362fs | 1 | 1,13 | 251108 | Pathogenic |
| gnomad0457 | c.439dupA | p.Thr147fs | 1 | 1,13 | 251122 | Likely pathogenic |
| gnomad0321 | c.817G>A | p.Ala273Thr | 1 | 1,13 | 251224 | Likely pathogenic |
| gnomad0256 | c.953A>T | p.His318Leu | 1 | 1,13 | 251230 | Likely pathogenic |
| gnomad0231 | c.1048C>T | p.Arg350Trp | 4 | 4,50 | 251258 | Likely pathogenic |
| gnomad0368 | c.703C>T | p.His235Tyr | 1 | 1,13 | 251270 | Likely pathogenic |
| gnomad0370 | c.700A>G | p.Thr234Ala | 3 | 3,38 | 251274 | Likely pathogenic |
| gnomad0374 | c.689A>G | p.Lys230Arg | 1 | 1,13 | 251276 | Likely pathogenic |
| gnomad0371 | c.698G>T | p.Arg233Leu | 1 | 1,13 | 251278 | Likely pathogenic |
| gnomad0240 | c.1000A>G | p.Ser334Gly | 1 | 1,13 | 251288 | Likely pathogenic |
| gnomad0241 | c.1000A>C | p.Ser334Arg | 1 | 1,13 | 251288 | Likely pathogenic |
| gnomad0235 | c.1027C>T | p.Arg343* | 1 | 1,13 | 251302 | Pathogenic |
| gnomad0237 | c.1020T>A | p.Asn340Lys | 3 | 3,38 | 251316 | Likely pathogenic |
| gnomad0438 | c.553_554insTG | p.Gln185fs | 2 | 2,25 | 251320 | Pathogenic |
| gnomad0599 | c.157G>A | p.Glu53Lys | 1 | 1,13 | 251350 | Likely pathogenic |
| gnomad0181 | c.1210G>T | p.Glu404* | 1 | 1,13 | 251364 | Pathogenic |
| gnomad0185 | c.1189G>A | p.Gly397Arg | 1 | 1,13 | 251386 | Likely pathogenic |
| gnomad0200 | c.1109-1G>A | NA | 1 | 1,13 | 251388 | Pathogenic |
| gnomad0097 | c.1362G>A | p.Met454Ile | 1 | 1,13 | 251400 | Likely pathogenic |
| gnomad0191 | c.1157A>C | p.Gln386Pro | 1 | 1,13 | 251404 | Likely pathogenic |
| gnomad0197 | c.1120C>T | p.Pro374Ser | 1 | 1,13 | 251408 | Likely pathogenic |
| gnomad0193 | c.1138dupA | p.Met380fs | 1 | 1,12 | 251412 | Likely pathogenic |
| gnomad0372 | c.698G>A | p.Arg233His | 5 | 5,00 | 282680 | Likely pathogenic |
| gnomad0447 | c.521C>G | p.Pro174Arg | 7 | 7,00 | 282696 | Pathogenic |
| gnomad0196 | c.1127A>C | p.Gln376Pro | 17 | 17,00 | 282834 | Likely pathogenic |
|  |  |  |  | **=0,000308** | **~ 1/3247** |  |

**Table S2 | Calculation of carrier frequency from the gnomAD database**

We filtered the variants retrieved from the gnomAD(36) database (v2.1.1 assessed on 2019-02-23), which we had annotated with InterVar(37) and manually curated ACMG(38) pathogenicity information from ClinVar and LOVD databases (Supplementary data file 2 sheet “FH_gnomAD”), for (likely) pathogenic variants. We then normalized the allele count (AC) to the maximum allele number in gnomAD and then divided the AC by the allele number to estimate the carrier frequency for (likely) pathogenic *FH* variants in this database. The result indicates that 1 in 3247 individuals (0.0308%) in gnomAD is a carrier.

| ID | HGVS_C | HGVS_P | AC | AC norm | Allele number | Reviewed ACMGclass |
| --- | --- | --- | --- | --- | --- | --- |
| bravo0065 | c.1391-2delA | NA | 2 | 2 | 125568 | Pathogenic |
| bravo0083 | c.1255T>C | p.Ser419Pro | 1 | 1 | 125568 | Likely pathogenic |
| bravo0135 | c.1189G>A | p.Gly397Arg | 2 | 2 | 125568 | Likely pathogenic |
| bravo0144 | c.1127A>C | p.Gln376Pro | 10 | 10 | 125568 | Likely pathogenic |
| bravo0147 | c.1093A>G | p.Ser365Gly | 1 | 1 | 125568 | Likely pathogenic |
| bravo0159 | c.1048C>T | p.Arg350Trp | 5 | 5 | 125568 | Likely pathogenic |
| bravo0161 | c.1020T>A | p.Asn340Lys | 1 | 1 | 125568 | Likely pathogenic |
| bravo0176 | c.923C>G | p.Ala308Gly | 2 | 2 | 125568 | Likely pathogenic |
| bravo0194 | c.817G>A | p.Ala273Thr | 2 | 2 | 125568 | Likely pathogenic |
| bravo0213 | c.736C>T | p.Gln246* | 1 | 1 | 125568 | Pathogenic |
| bravo0217 | c.700A>G | p.Thr234Ala | 1 | 1 | 125568 | Likely pathogenic |
| bravo0218 | c.698G>A | p.Arg233His | 4 | 4 | 125568 | Likely pathogenic |
| bravo0219 | c.697C>T | p.Arg233Cys | 2 | 2 | 125568 | Likely pathogenic |
| bravo0264 | c.555+1G>C | NA | 2 | 2 | 125568 | Pathogenic |
| bravo0270 | c.521C>G | p.Pro174Arg | 6 | 6 | 125568 | Pathogenic |
| bravo0272 | c.504delA | p.Glu168fs | 1 | 1 | 125568 | Likely pathogenic |
| bravo0275 | c.473G>A | p.Ser158Asn | 1 | 1 | 125568 | Likely pathogenic |
| bravo0313 | c.301C>T | p.Arg101* | 2 | 2 | 125568 | Pathogenic |
| bravo0414 | c.65T>A | p.Leu22* | 1 | 1 | 125568 | Likely pathogenic |
| bravo0423 | c.40dupC | p.Leu14fs | 2 | 2 | 125568 | Likely pathogenic |
|  |  |  |  | **= 0,000390** | **~ 1/2563** |  |

**Table S3 | Calculation of carrier frequency from the BRAVO database**

We filtered the variants retrieved from the BRAVO (Freeze5 assessed on 2019-02-23) database, which we had annotated with InterVar and manually curated pathogenicity information from ClinVar and LOVD databases (Supplementary data file 2 sheet “FH_BRAVO”), for (likely) pathogenic variants and divided the allele count (AC) by the allele number to estimate the carrier frequency for (likely) pathogenic *FH* variants in this database. The result indicates that 1 in 2563 individuals (0.0390%) in BRAVO is a carrier.
